# Stem cell morphology defines functional heterogeneity and therapeutic vulnerabilities in glioblastoma

**DOI:** 10.1101/2025.03.24.644884

**Authors:** Carlotta Barelli, Matteo Bonfanti, Filippo Mirabella, Dario Ricca, Negin Alizadehmohajer, Roberta Bosotti, Ilaria Bertani, Viktoria Sokolova, Alberto Campione, Giovanni M. Sicuri, Clelia Peano, Stefania Faletti, Roberto Stefini, Nereo Kalebic

**Author notes:** Equal contribution (Matteo Bonfanti, Filippo Mirabella).

## Abstract

Glioblastoma (GBM) is an aggressive brain tumor and an unmet clinical need due to its invasiveness and therapy-resistance. These features are driven by glioblastoma stem cells (GSCs), which exhibit remarkable functional heterogeneity. However, GSC transcriptional profiling alone cannot predict clinically relevant behaviors. Here, we developed CellShape-seq, a spatial transcriptomics platform that integrates cell morphology with transcriptome. This identified three GSC morphoclasses corresponding to distinct transcriptomic states and functions: (1) nonpolar cells show differentiation and therapy sensitivity, (2) elongated cells are invasive, and (3) multipolar cells form intercellular networks. Importantly, chemoresistance is morphoclass-specific: elongated GSCs depend on YAP/TEAD1 signaling, while multipolar GSCs rely on gap junction–mediated networks. Targeting these vulnerabilities with specific inhibitors sensitized resistant GSC morphoclasses to temozolomide (TMZ) in patient-derived organoids. Our findings demonstrate that morphology provides critical insights into GSC behavior and establish a rationale for morphology-informed therapies to overcome resistance and improve outcomes in GBM.

**Graphical abstract:** 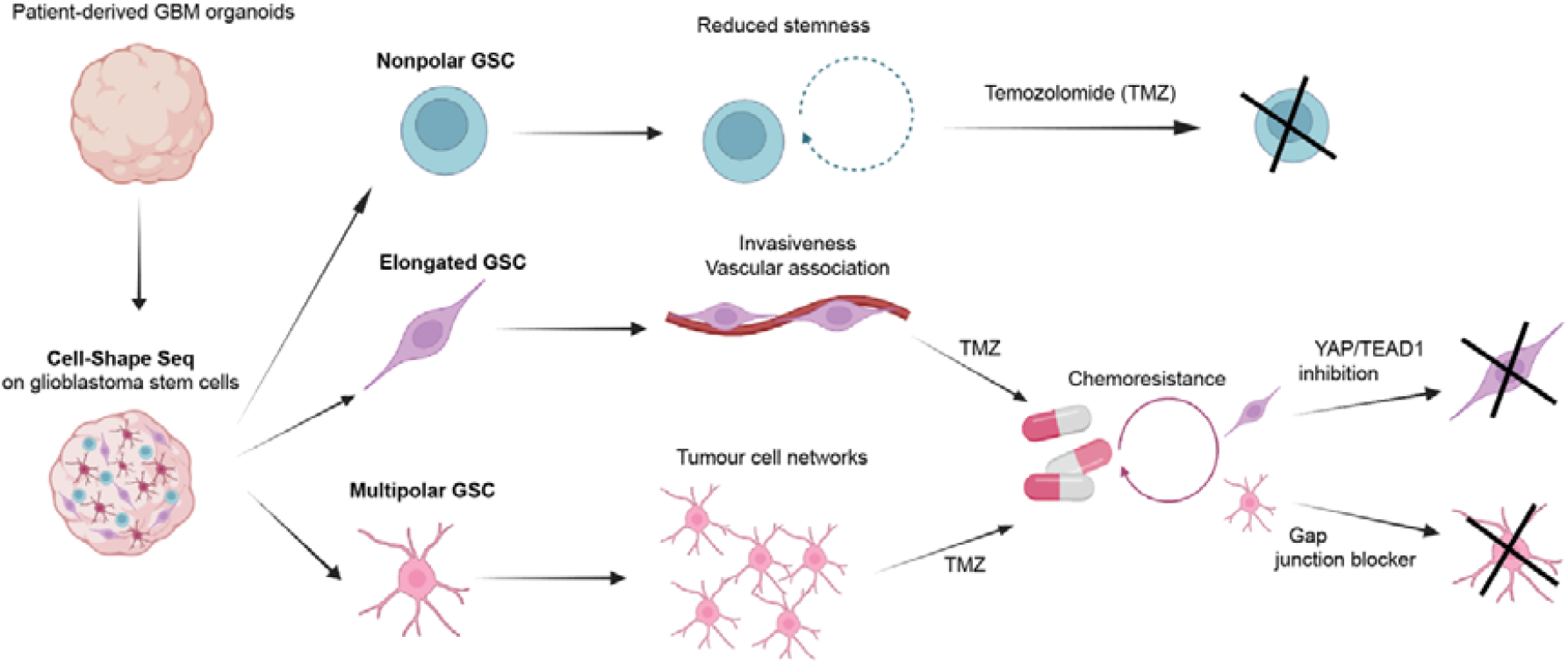

## Introduction

Glioblastoma (GBM) is the most common and aggressive brain tumor, representing an unmet clinical need, with a median survival rate of only 15 months (Tran and Rosenthal, 2010). Characterized by extensive intra- and inter-tumoral heterogeneity, resistance to conventional therapies, and aggressive infiltration into healthy brain tissue, GBM almost inevitably relapses (Marenco-Hillembrand et al., 2020; Stupp et al., 2005). Central to these features are GBM stem cells (GSCs), a subpopulation characterized by remarkable heterogeneity (Neftel et al., 2019) which drives tumor invasion and chemoresistance (Gimple et al., 2019; Liu et al., 2006; Xie et al., 2014; Xie et al., 2022). However, mechanisms underlying GSC heterogeneity remain unclear. Scrutinizing GSC biology and elucidating the mechanisms driving their heterogeneity is hence required for developing novel therapeutical approaches tackling GBM aggressiveness.

Transcriptional profiling has been valuable to extract the heterogeneity of cellular states in GBM and GSCs (Garofano et al., 2021; Neftel et al., 2019; Wang et al., 2017; Yu et al., 2020). However, the transcriptome alone cannot fully account for the heterogeneity in GSC behavior and is therefore insufficient to link clinically relevant functions to specific GSC states and ultimately to predict patient prognosis. In this regard the cell morphology has been proven to be a complementary, reliable predictor of cell behavior in various oncological contexts (Alizadeh et al., 2020; Conner et al., 2024; Sali et al., 2024). Interestingly, GSCs exhibit morphological stability, as we have recently showed that the morphotype identity is both inherited from mother to daughter cells and maintained throughout the interphase (Barelli et al., 2025). Moreover, GBM cells exhibit specialized cellular protrusions connecting tumor cells into a multicellular network that enhances tumor resilience and repair capacity (Osswald et al., 2015; Pinto et al., 2020). This provides a basis to link specific cell morphologies with distinct functions in GBM.

GSCs share striking transcriptional similarities with neural progenitor cells, particularly basal or outer radial glia (bRG) (Azzarelli et al., 2018; Bhaduri et al., 2020; Couturier et al., 2020; Neftel et al., 2019). Interestingly, cell morphology is a critical feature for key bRG functions, such as proliferation and support for neuronal migration (Kalebic and Huttner, 2020). In fact, we recently observed that GSCs exhibit morphological heterogeneity, with morphotypes reminiscent of those seen in embryonic bRGs (Barelli et al., 2025). Furthermore, both GSCs and bRGs share specific morphoregulatory proteins, such as ADD3 (Barelli et al., 2025; Kalebic et al., 2019), highlighting potential mechanistic links. This prompts us to investigate if GSCs’ morphology, combined with their specific transcriptomic signature, could be linked to GSC heterogeneous behavior.

While recent studies have suggested that morphological traits could predict functional behaviors in various cancer cell lines (Alizadeh et al., 2020; Barker et al., 2022; Conner et al., 2024; Da et al., 2022; Sali et al., 2024; Wu et al., 2020), these analyses were rarely focused on stem cells. In addition, these results predominantly relied on 2D cultures, which fail to fully recapitulate the complexities of in vivo environments. Of note, also in the context of neurodevelopment, the bRG morphology regulates cellular functions by modulating interaction with the microenvironment (Kalebic and Huttner, 2020). In this study we employed patient-derived organoids and assembloids, which contain both the cellular heterogeneity and the extracellular signals critical for the regulation of cell morphology and allow us to study how GSC morphology shapes the interactions with the surrounding cells. We further developed a customized spatial biology approach linking cell morphology with the transcriptome (CellShape-seq), which allowed us to identify three different GSC morphoclasses (nonpolar, elongated and multipolar) with distinct cell states. Our functional assays show that different GSC morphotypes convey distinct clinically relevant functions and therapeutic vulnerabilities. Furthermore, our morphotype-specific pharmacological treatments strikingly reduced GBM chemoresistance, highlighting that cell morphology is a critical aspect of GBM heterogeneity and a promising target for novel cell biology-informed therapies.

## Results

### GSCs are morphologically heterogeneous in patient-derived organoids

We generated patient-derived GBM organoids using a protocol which maintains the original 3D tumor architecture and native microenvironment without dissociation (Jacob et al., 2020) (fig. S1A, Fig. 1A, B). Organoid viability was confirmed by markers for mitotic and actively cycling cells and lack of the apoptotic signals (fig. S1B, C). Using a panel of markers, we validated the presence of GSCs, along with neurons, astrocytes and fragments of blood vessels in GBM organoids (Fig. 1B, fig. S1D-G). For most future purposes, GSCs were identified as cells expressing both SOX2 and nestin (Fig. 1A, B), with the nestin signal used to track cell shape. We hence identified six distinct GSC morphotypes, defined as categories of cells with specific and consistent morphological features (Fig. 1C). We further grouped these morphotypes into three main morphoclasses (families of morphotypes with similar core features), comparably to what we previously established in 2D cancer cell lines (Barelli et al., 2025). The three morphoclasses were: nonpolar (encompassing both flat and round nonpolar cells and accounting for ∼20% of all GSCs), elongated (comprising radial or monopolar and bipolar cells; ∼30% of total) and multipolar (containing both flat and round nonpolar cells; ∼50% of total) (Fig. 1C, D). Importantly, organoids faithfully recapitulated the morphological distribution of GSCs observed in patient samples for at least 15 weeks (Fig. 1D, fig. S1H-J), consistently with what previously reported for gene expression (Jacob et al., 2020). Thus, GBM organoids recapitulate and maintain the morphological heterogeneity of GSCs, thereby providing a robust model to study its functional implications.

**Figure 1.**
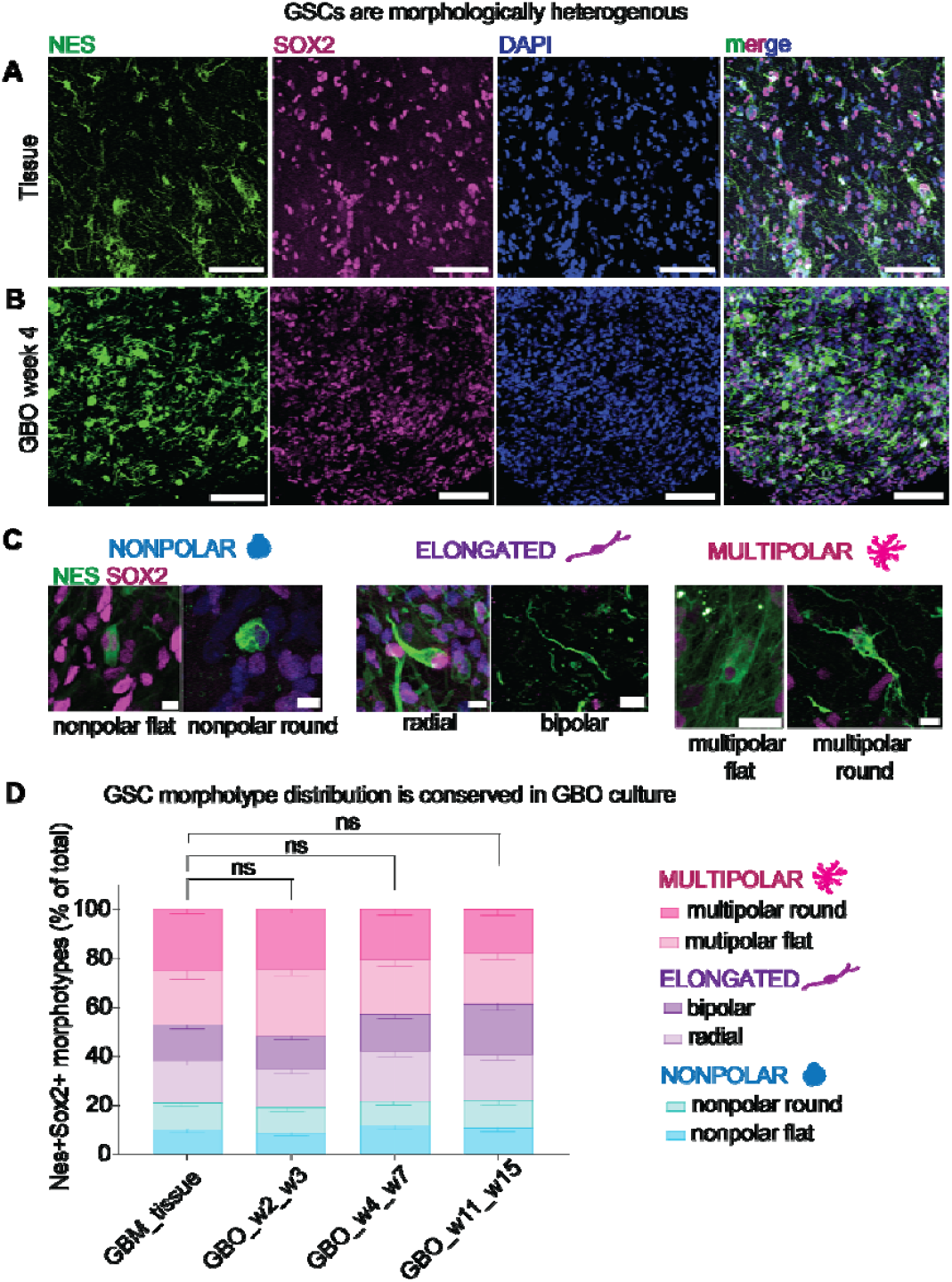
GSCs are morphologically heterogenous in patient-derived tissue and organoids. **(A-B)** Representative microscopy images of Nes+ SOX2+ GSCs exhibiting different morphologies in patient primary tissue (A) and patient-derived glioblastoma organoid (GBO, B). Scale bars, 100 µm. **(C)** Representative microscopy images of the six Nes+ SOX2+ GSC morphotypes grouped into three morphoclasses. Nonpolar cells comprise nonpolar flat and nonpolar round; elongated comprise radial and bipolar; multipolar comprise multipolar flat and round. Scale bars, 10 µm. **(D)** Quantitative analysis of the distribution of GSC morphotypes and morphoclasses in tissue and GBM organoids (GBO) at different culture timings. Morphotype distribution is not altered upon organoid culture. Error bars, SEM; Two-way ANOVA with Sidak post-hoc test; N=4 patients (P01, P02, P03, P04), 3+ organoids per patients, 3 fields of view per organoid/tissue section.

### CellShape-seq – a cell morphology based spatial transcriptomics approach – shows that GSC morphotypes correspond to distinct transcriptomic states

To transcriptionally characterize individual morphoclasses, we performed spatial transcriptomics. While this technique has been valuable at integrating tissue morphology with cell identity and function (Bao et al., 2022; Song et al., 2024; Walker et al., 2022), it has not been explored to utilize morphology of individual cells. We hence developed CellShape-seq, a custom spatial transcriptomics approach by adapting the GeoMX platform. Our approach consists of manual segmentation of individual cells belonging to different morphoclasses, followed by sequencing-based morphoclass-specific whole transcriptome analysis from multiple patient-derived organoids (Fig. 2A, B). This analysis revealed patient-dependent clustering (fig. S2A), consistent with previous reports in GBM (Couturier et al., 2020; Garofano et al., 2021). To account for this, we performed pathway deconvolution analysis as done previously (Garofano et al., 2021). This demonstrated that nonpolar GSCs are transcriptionally more distant from elongated and multipolar GSCs (Fig. 2C, D, fig. S2B). Clustering with the Leiden algorithm further supported this separation, placing nonpolar cells apart from elongated and multipolar populations (Fig. 2E, F).

**Figure 2.**
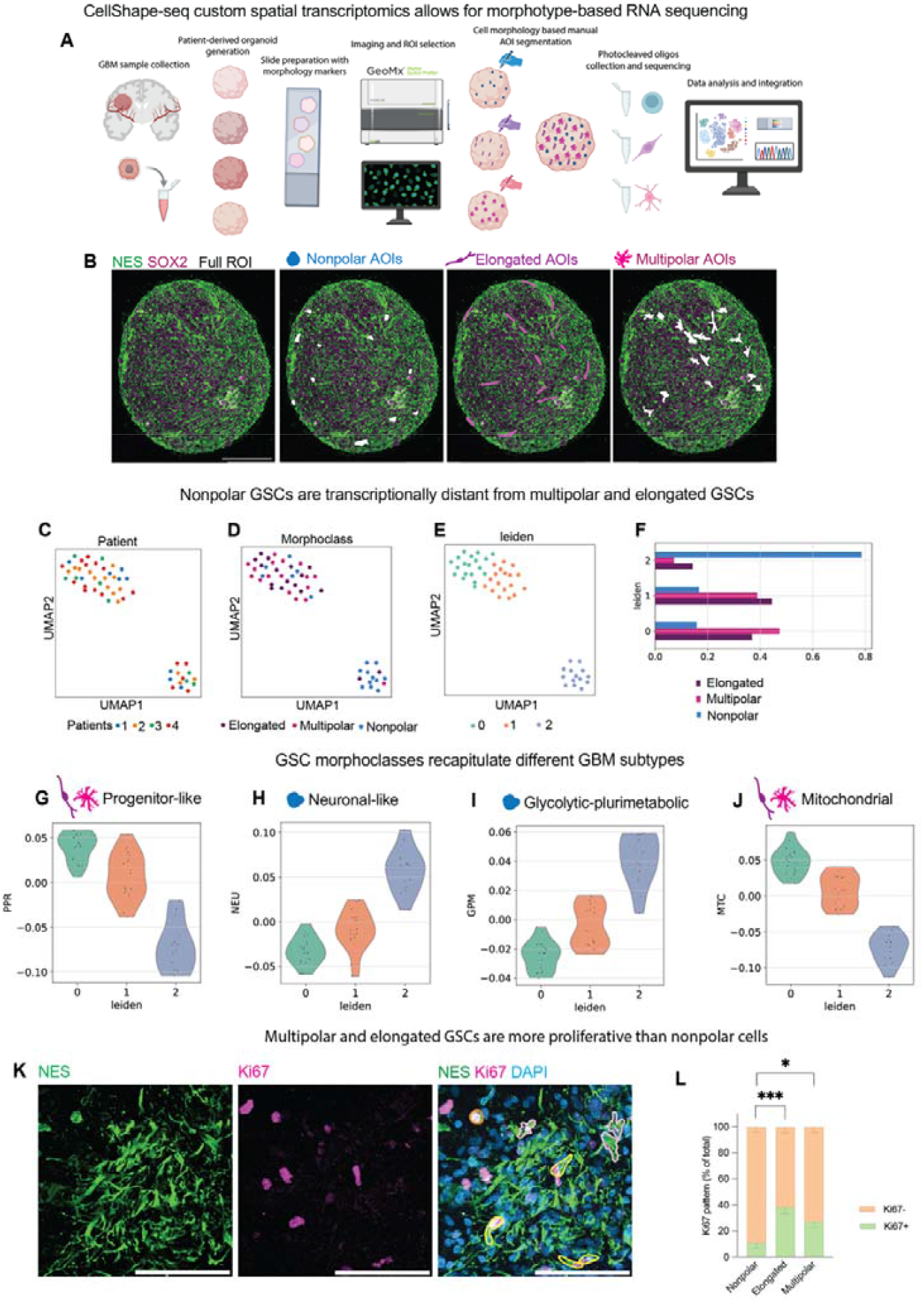
CellShape-seq reveals that GSCs have morphoclass-specific transcriptomic signatures. **(A, B)** CellShape-seq is a custom spatial transcriptomics approach to select cells based on their individual morphologies. (A) Schematics of the pipeline. Following the collection of primary GBM samples, patient-derived organoids are generated, sliced, stained and hybridized with probes. Upon imaging on GeoMX DSP individual cell morphologies are segmented and photocleaved oligos are collected from the regions of interest (ROIs), followed by data analysis. (B) Representative image of a GBM organoid (ROI) stained with Nestin and SOX2 and acquired with the GeoMX DSP to identify the Areas of illumination (AOIs, cell segmentations of the same morphoclass). AOI segmentations for nonpolar, elongated and multipolar GSCs. Scale bars, 500 µm. **(C-F)** Clustering of the pathway deconvolution analysis results. Uniform Manifold Approximation and Projection (UMAP) plot of pathway deconvolution results for each AOI, computed as described in the Methods section. The three panels (C-E) show coloring by patient (P03, P04, P05, P06) (C), morphoclass (D), and clusters computed with Leiden algorithm (E). The accompanying bar plot (F) displays the number of AOIs per cell type within each cluster. **(G-J)** Functional scores of the pathway deconvolution clusters. Violin plots depicting different functional scores— Progenitor-like (G), Neuronal-like (H), Glycolytic-plurimetabolic (I), and Mitochondrial (J)— grouped by Leiden clusters. Functional scores for each AOI were computed by averaging pathway enrichment values over the pathway classes described in (Garofano et al., 2021). **(K-L)** Nonpolar GSCs are less proliferative than elongated and multipolar cells. (K) Representative microscopy image of primary GBM tissue stained with Nestin (green) and Ki67 (magenta). Cell outlines, yellow (elongated), pink (multipolar), orange (nonpolar). Scale bars, 100 µm. (L) Quantitative distribution of Ki67+ and Ki67-GSCs in each morphoclass. Error bars, SEM. N= 6 patients (P01, P02, P03, P12, P18, P21). Two-way ANOVA with Tukey post-hoc test. Elongated vs nonpolar, ***, P<0,001; multipolar vs nonpolar, *, P<0,05.

Such results prompted us to explore potential resemblance of Leiden clusters with four GBM subtypes, which embody developmental and metabolic attributes, as described previously (Garofano et al., 2021). Functional scoring of pathway enrichment values linked nonpolar GSCs to neuronal-like GBM subtype and elongated and multipolar cells to progenitor-like subtype (Fig. 2G, H). In the context of the metabolic states, elongated and multipolar cells were associated to a GBM subtype dependent on oxidative phosphorylation (termed mitochondrial GBM state (Garofano et al., 2021)), whereas nonpolar GSCs showed more resemblance to the glycolytic-plurimetabolic subtype (Fig. 2I, J). Such metabolic signature of nonpolar GSCs was driven by the pathways related to differentiation, inflammation and hormonal metabolism (fig. S2B, S3A) and not by signatures of metabolic plasticity associated with GBM aggressiveness (fig. S3B). As nonpolar GSCs were linked to a neuronal-like GBM cell state and were enriched in differentiation pathways, we hypothesized differences in proliferative index across GSC morphoclasses. Indeed, our Ki67 analysis (Fig. 2K, L) showed increased proliferation of elongated and multipolar GSCs compared to nonpolar GSC, in agreement with the transcriptomic data. Next, we performed differential expression analysis of nonpolar GSCs *vs* the rest (fig. S3C) and further detected reduced expression of both protein translation genes, potentially linked to their smaller size, and genes involved in oxidative phosphorylation, in line with analysis of GBM subtypes (fig. S3D). Overall, nonpolar GSCs display decreased stemness compared to elongated and multipolar cells, as well as decreased aerobic metabolism and cell size.

Overall, CellShape-seq enabled the creation of the first morphology-based atlas of GSCs in primary patient samples. Surprisingly, we found that GSC morphoclasses align with distinct transcriptomic states reflective of clinically relevant GBM subtype distributions. Notably, elongated and multipolar GSCs showed the strongest resemblance to a progenitor-like GBM state, exhibited higher Ki67 expression, and displayed more complex morphologies. These features prompted us to functionally investigate these two subpopulations.

### Elongated GSCs are invasive and associated to blood vessels

Differential gene expression analysis comparing elongated GSCs to other morphoclasses revealed significant enrichment in genes related to cell migration and adhesion (Fig. 3A, fig. S4A, B). Performing a functional gene pathway analysis, we found that amongst the top 10 modulated genes seven were related to neurodevelopment, axon growth and tumor cell migration (fig. S4C, D). We put these results in the context of previous studies suggesting that cell elongation can predict invasiveness across various cancer cell types (Baskaran et al., 2020; Fayzullin et al., 2016). Among the genes enriched in elongated GSCs, we detected CD99 (fig. S5A, B), previously implicated in GBM invasiveness and cell elongation (Cardoso et al., 2019; Seol et al., 2012), and observed an enrichment of CD99+ cells along the blood vessels in three out of five patient tumor samples that we examined (fig. S5C-F).

**Figure 3.**
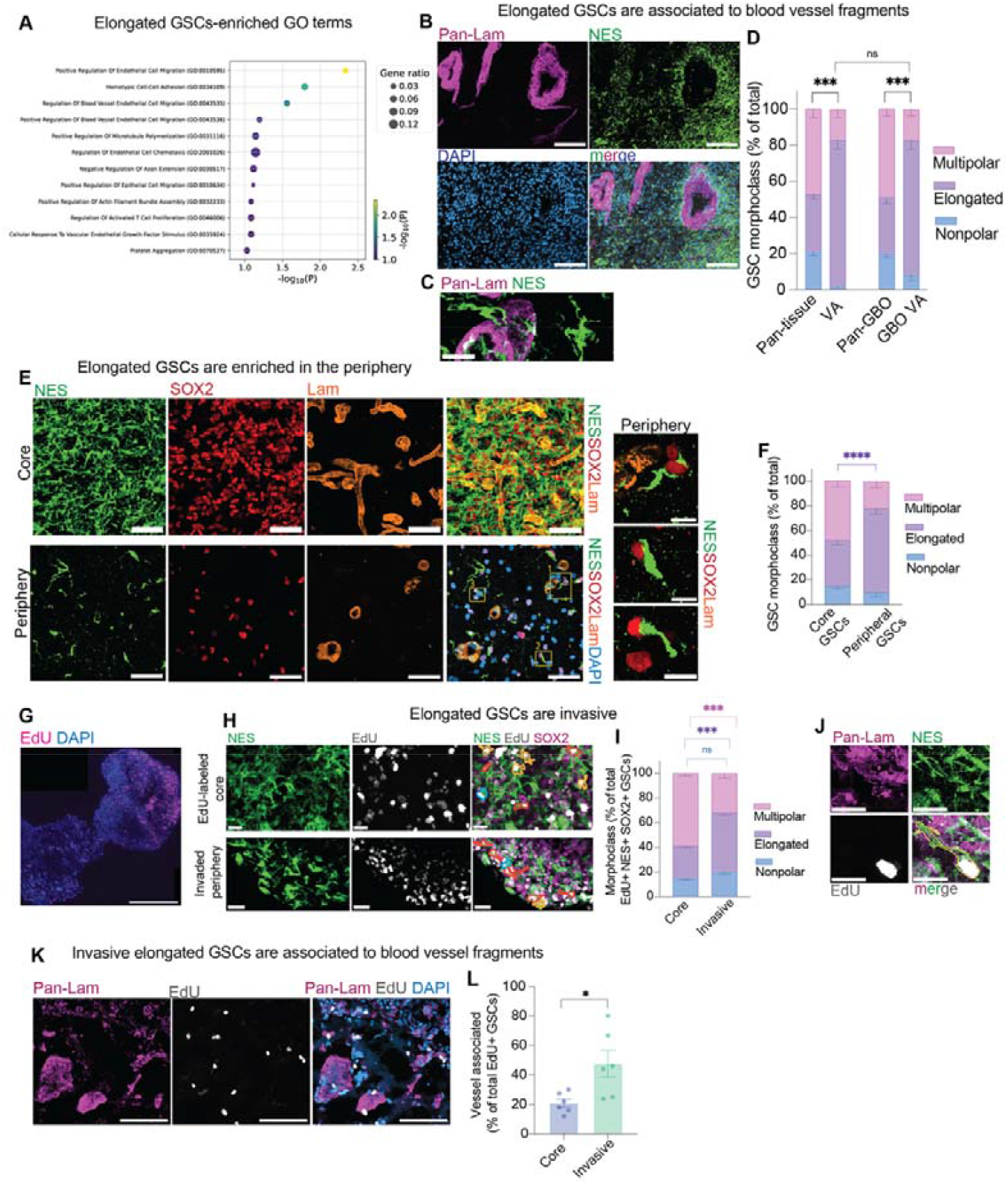
Elongated GSCs are invasive and associated to blood vessels. **(A)** Enrichment of selected Gene Ontology (GO) Biological Process (BP) terms in elongated cells. In the plot, the x-axis position and dot color represent the significance level of the pathway enrichment, while the dot size indicates the proportion of genes associated with each GO term that are differentially expressed in our analysis. Just some selected GO terms which we functionally validated were selected (view full graph in fig. S4B). **(B-D)** Elongated GSCs are associated to blood vessel fragments. Representative overview (B) and high magnification view (C) of Nes+ GSCs in the parenchyma and associated to Laminin+ blood vessel fragments. Scale bars, 100 µm (B), 20 µm (C). (D) Quantitative distribution of the three morphoclasses associated to the blood vessel fragments compared to the overall distribution in tissue and GBM organoids (GBO). Error bars; SEM. Two-way ANOVA with Sidak post-hoc test. ****, P<0,0001. N=4 patients (P01, P03, P12, P17). **(E-F)** Elongated GSCs are enriched in the periphery of the primary tumor. (E) Representative microscopy image of core (top) and peripheral (bottom) patient GBM tissue stained with Nestin (green), SOX2 (red), Laminin (orange) and DAPI (blue, in the merge). Elongated GSCs are enriched in the periphery (see the three close ups). Scale bars, 100 µm (F) Quantitative distribution of the three morphoclasses in patient tissue sections of the core and periphery of the tumor. Error bars, SEM. N=5 patients (P01, P03, P12, P15, P17). Two-way ANOVA with Sidak multiple comparisons; ****, P<0,0001. **(G-I)** Elongated GSCs are invasive. (G) Representative microscopy image of one assembloid with EdU+ (magenta) core organoid fused with a peripheral organoid invaded by EdU+ GSCs. Scale bar, 500 µm. (H) Nes+ SOX2+ EdU+ cells in the core and peripheral part of the assembloid have different morphologies (cell segmentations in colors: red-elongated, blue-nonpolar; orange-multipolar). Scale bars, 20 µm. (I) Quantitative distribution of EdU+ Nes+ SOX2+ morphoclasses in the core and in the invaded part of the assembloid. Error bars, SEM. N= 21 assembloids from 5 patients (P07, P08, P11, P12, P15). Two-way ANOVA with Sidak multiple comparisons; ****, P<0,0001. **(J-L)** Invading elongated GSCs are associated to blood vessels. (J) Representative microscopy image of an elongated invasive EdU+ GSC associated to a vessel. Scale bar, 20 µm. (K) Representative microscopy image of EdU+ invading cells associated to Laminin+ vessels. Scale bar, 100 µm. (L) Quantification of EdU+ GSCs associated to the vessels in the core and in the invaded part of the assembloid. N=6 assembloids from 3 different patients (P08, P12, P15). Error bars, SEM. Student’s t-test; *, P= 0,0189.

Given that invading GBM cells frequently use vasculature as migratory paths (Baker et al., 2014; Krusche et al., 2016; Monzo et al., 2016; Watkins et al., 2014), we investigated the relationship between elongated GSCs and blood vessel fragments in our culture systems. Microscopy analysis showed that elongated cells account for around 80% of vessel-attached (VA) GSCs both in GBM tissue and organoids, whereas they comprise only around 30% of total GSCs (Fig. 3B-D), indicating a strong enrichment of elongated GSCs around blood vessels. Given the potential role of elongated GSCs in invasiveness we compared the GSC morphology in the tumor core with the periphery of the primary GBM samples. Spatial analysis by immunofluorescence showed a remarkable enrichment of elongated GSCs in the GBM periphery (Fig. 3E, F), suggesting that elongated GSC may be implicated in tumor invasiveness. Next, we further examined the tumor periphery by distinguishing invasive and perivascular niches (Barthel et al., 2022; Hambardzumyan and Bergers, 2015). We identified the invasive niche through enrichment for neuronal, microglial and metalloproteinase markers (fig. S6A, B) and the perivascular niche by presence of pericytes and perivascular macrophages (fig. S6C). Analysis of the morphoclass distribution showed a striking enrichment of elongated GSCs in both the invasive and perivascular niches compared to core tumor (fig. S6D). Together, these data suggest a strong spatial enrichment of elongated cells in tumor regions implicated in invasiveness.

Hence, we functionally assessed the involvement of elongated GSCs in GBM invasiveness using invasion assays in assembloids. While previous assembloid models mostly fuse GBM organoids with developing cortical organoids lacking vasculature (da Silva et al., 2018; Kim et al., 2025; Liang et al., 2022; Roth et al., 2023), we fused them with peripheral tumor organoids, which do contain fragments of blood vessels. We labeled GBM cells in the core with a single pulse of EdU and examined their invasion into the peripheral organoid ten days post fusion (Fig. 3G, H, fig. S6E). The relative distribution of morphoclasses among EdU+ cells in the core organoid was comparable to untreated controls (compare Fig. 3I core and 4D pan-GBO), indicating no bias in S-phase progression across morphoclasses. Strikingly, despite representing ∼30% of GSCs in the core, the elongated EdU+ GSCs accounted for ∼50% of cells invading into the peripheral organoid (Fig. 3I). Furthermore, vessel-associated EdU+ GSCs doubled in proportion in the invaded periphery compared to the core (Fig. 3J-L).

These functional assays in assembloids strongly support that elongated GSCs drive invasion along blood vessels, consistent with their transcriptional profile and localization within the tumor niches. Collectively, our data show that elongated cells associate to vessels and are highly invasive.

### Multipolar GSCs form functional networks

Transcriptomic analysis revealed that multipolar GSCs exhibit enrichment in genes related to morphoregulation, growth of cell protrusions and calcium signaling (Fig. 4A, fig. S7A, B). Performing a functional gene pathway analysis, we observed that the top 10 modulated genes in multipolar GSCs relate largely to cytoskeletal dynamic, protrusion growth and cell metabolism (fig. S7C, D). We link these data with the notion that GBM cells form a network of inter-cellular connections that are important for maintaining the aggressive and chemo-resistant nature of the tumor (Barelli et al., 2025; Hai et al., 2024; Pinto et al., 2020; Venkataramani et al., 2022a). Such connections, mediated by tumor microtubes and tunneling nanotubes, enable inter-cellular communication through calcium signaling (Hausmann et al., 2023; Osswald et al., 2015; Smith et al., 2011; Vargas et al., 2019) and mitochondrial exchange, similar to calcium waves in radial glia during neurodevelopment (Rash et al., 2018; Weissman et al., 2004). We hence sought to examine the relative contribution of multipolar cells to such networks in GBM organoids.

**Figure 4.**
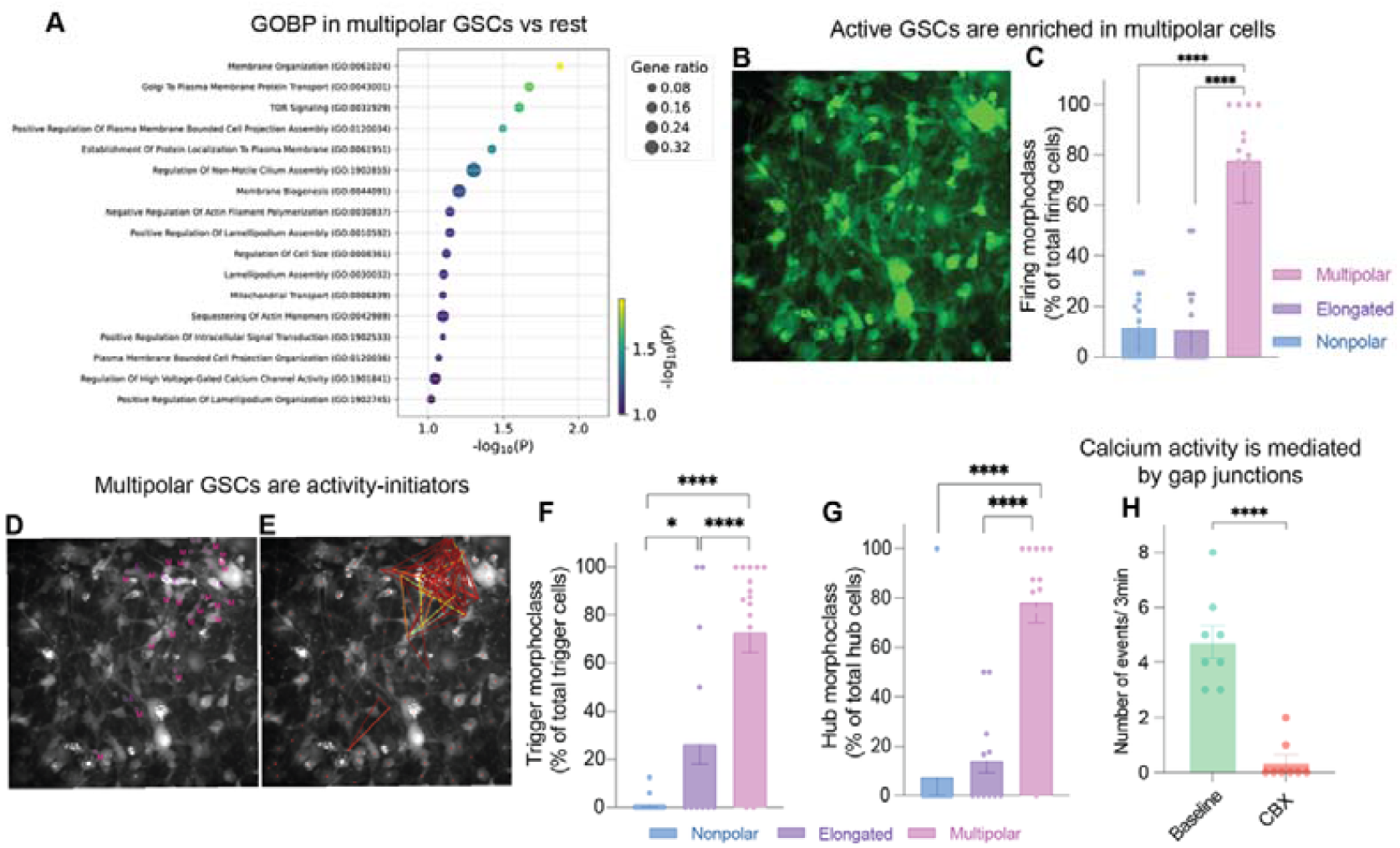
Multipolar GSCs form a functional resistant network. **(A)** Enrichment of selected GO Biological Process terms in multipolar cells. In the plot, the x-axis position and dot color represent the significance level of the pathway enrichment, while the dot size indicates the proportion of genes associated with each GO term that are differentially expressed in our analysis. Just some selected GO terms which we functionally validated were selected (view full graph in fig. S8B). **(B-C)** Multipolar cells are the most active. (B) Max intensity projection (MIP) of the live Fluo4 calcium imaging movie (see Movie S2). Frame width, 389 µm. (C) Quantification of firing morphoclasses over total firing cells. Error bars, SEM. One-way ANOVA with Tukey post-hoc test; ****, P<0,0001. N=39 organoids from 7 different patients (P07, P08, P10, P11, P12, P13, P14). **(D-G)** Multipolar cells are activity-initiators. (D) MIP of the live Fluo4 calcium imaging movie with morphoclass label in active cells (magenta, multipolar (M); purple, elongated (E)) (see Movie S2). (E) Example of connectivity map starting from a hub cell and spreading to neighboring cells. Frame width, 389 µm. (F, G) Quantitative distribution of trigger morphoclasses over total trigger cells (F) and of hub morphoclasses over total hub cells (G). Error bars, SEM. One-way ANOVA with Tukey post-hoc test; *, P<0,05; ****, P<0,0001. N=same as (C). **(H)** Calcium activity is mediated by gap junctions. Number of calcium events within 3 min recordings at baseline and after 15 min of treatment with 100 µM carbenoxolone (CBX). N= 7 organoids from 3 patients (P10, P14, P15). Error bars, SEM. Student’s t-test, ****, P<0,0001.

We performed live calcium imaging in GBM organoids using Fluo4 sensor (Movies S1-S3) and analyzed the morphology of the active cells (Fig. 4B). Remarkably, almost 80% of the active cells were multipolar (Fig. 4C and compared to fig. S8A, showing overall distribution of morphotypes in the same samples). Previous study (Hausmann et al., 2023) identified GBM cells displaying rhythmic calcium oscillations that trigger calcium waves within tumor cell networks (trigger cells, see Movies S1 and S3), serving as hubs for intercellular communication (hub cells, see Movie S2). By mapping calcium connectivity, we observed that multipolar cells predominantly act as both trigger and hub cells, further emphasizing their fundamental role in GBM networks (Fig. 4D-G). Such connections between GBM cells are often mediated through gap junctions (Osswald et al., 2015; Sinyuk et al., 2018). To examine this, we treated the organoids with the gap junction blocker Carbenoxolone (CBX) and we observed a striking reduction in calcium activity (Fig. 4H and compare Movies S3 and S4). We further performed immunostaining for connexin-43 (Cx43) and observed that multipolar GSCs in GBM organoids are connected through Cx43+ gap junctions (fig. S8B). Collectively, our calcium imaging in organoids shows that multipolar GSCs establish a gap junction-mediated functional intercellular network.

### GSCs exhibit morphoclass-specific therapeutic vulnerabilities

Multicellular networks in GBM, formed via gap junction-mediated communication, have been strongly implicated in therapeutic resistance (Hekmatshoar et al., 2018; Kolba et al., 2019; Osswald et al., 2015; Potthoff et al., 2019; Wang et al., 2018; Weil et al., 2017). Previously, we demonstrated that the morphoregulatory protein adducin-3 (ADD3) promotes complex GSC morphology, intercellular connectivity, and resistance to temozolomide (TMZ), the standard chemotherapeutic for GBM (Barelli et al., 2025). To further explore how morphology impacts chemoresistance, we treated GBM stem cells with TMZ and observed that surviving, TMZ-resistant, cells predominantly displayed multipolar and elongated morphologies, whereas TMZ-sensitive cells exhibited simpler flat shapes without protrusions (fig. S9A-C). We next treated patient-derived GBM organoids with TMZ and detected an enrichment of multipolar and elongated GSCs at the expense of nonpolar cells after 10 days of metronomic treatment (Fig. 5A, B, fig. S9D). These results strongly support a link between morphological complexity and chemoresistance in GSCs.

**Figure 5.**
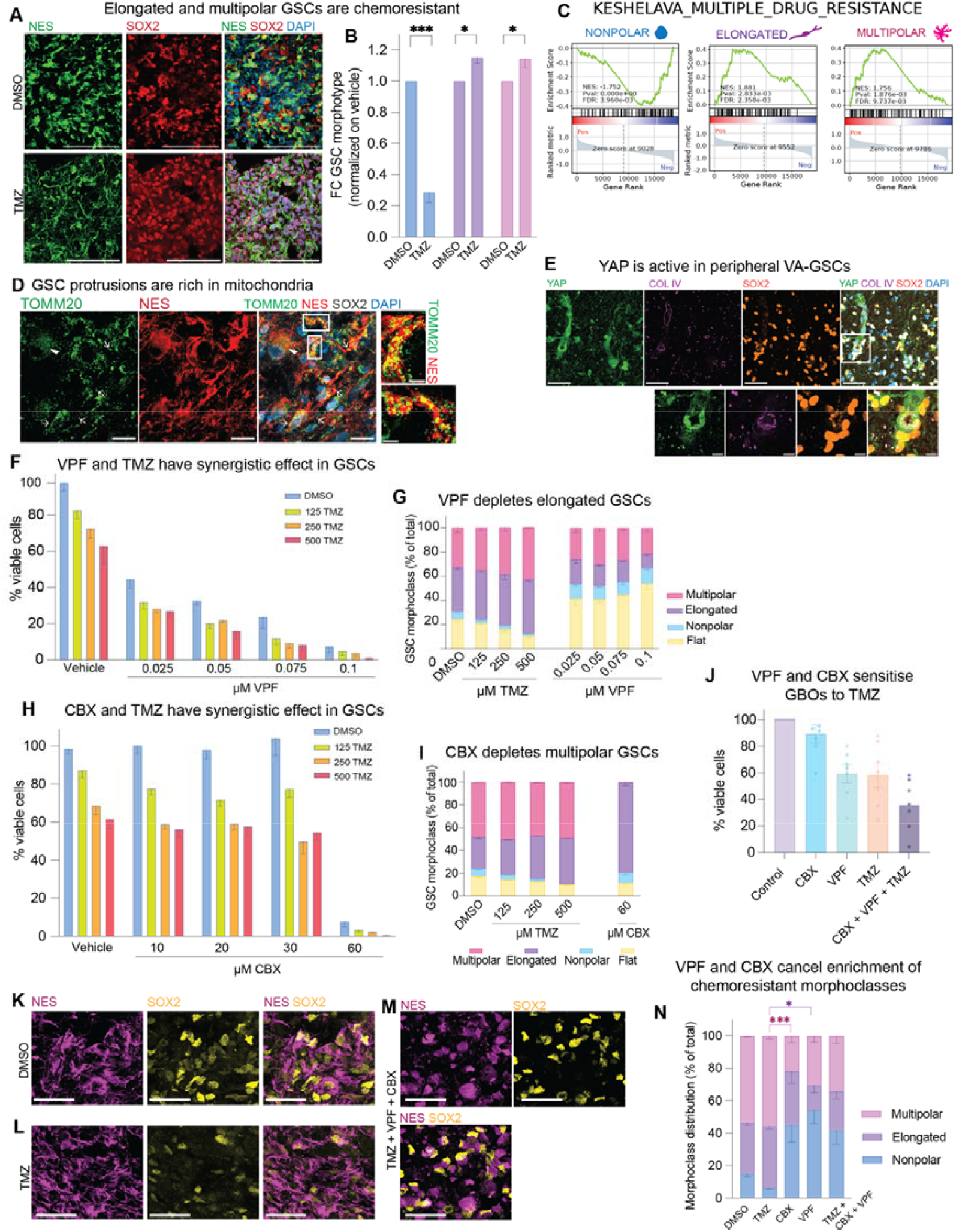
GSCs have morphoclass-specific therapeutic vulnerabilities. **(A, B)** Multipolar and elongated GSCs are resistant to temozolomide (TMZ). (A) Representative microscopy image of GBM organoids (GBO) treated with 500 µM TMZ for 10 days (bottom) or DMSO control (top) stained with Nestin (green), SOX2 (red) and DAPI (blue). Scale bars, 100 µm. (B) Quantification of (A). Data are expressed as fold change and normalized on DMSO (set to 1 for each morphoclass). Error bars, SEM. Two-way ANOVA with Sidak post-hoc test; ***, P<0,001, *, P<0,05. N=3 experiments with 4 different patients (P10, P16, P18, P20). **(C)** Gene set enrichment analysis (GSEA) plot of the human gene set (Keshelava et al., 2007) comprising genes that are overexpressed in multidrug-resistant neuroblastoma cell lines. Normalized enrichment score (NES), P-value (Pval) and false discovery rate (FDR) are indicated in the plots for each of the three morphoclasses. (D) GSC protrusions contain mitochondria. Representative microscopy images of Nestin+ SOX2+ GSCs containing mitochondria (revealed by TOMM20 staining, green) in their protrusions in human GBM sample. Arrows, Multipolar GSCs’ protrusions rich in big mitochondria; Arrowheads, flat non-polar GSC with smaller mitochondria. Insets, two examples of GSC protrusions rich in mitochondria. Scale bars, 20 µm. **(E)** YAP is present in the nucleus and cytoplasm of peripheral vessel-associated (VA)-GSCs. Representative microscopy image of tumor periphery and insets with an example of the blood vessel fragment with YAP+ VA-GSCs. Scale bars, 50 µm (top) and 10 µm (bottom, inset). **(F)** VPF and TMZ have synergistic effects. GSC viability after 96h of 0.025-0.1 µM VPF treatment alone (blue) or in combination with 125 µM TMZ (green), 250 µM TMZ (orange), 500 µM TMZ (red). Measurements are normalized on DMSO control (100%). N= 1 GSC line (P03); 3 biological replicates. Error bars, SEM. See Fig. S11 with the corresponding microscopy images along with the quantifications for the second cell line. **(G)** VPF depletes elongated GSCs. GSC morphotype distribution after 96h treatment with 125-500 µM TMZ and 0.025-0.1 µM VPF. N= 1 GSC line (P03); 3 biological replicates. Error bars, SEM. **(H)** CBX and TMZ have synergistic effects. GSC viability after 96h of 10-60 µM CBX treatment alone (blue) or in combination with 125 µM TMZ (green), 250 µM TMZ (orange), 500 µM TMZ (red). Measurements are normalized on DMSO control (100%). N= 1 GSC line (P06); 3 independent replicates. Error bars, SEM. **(I)** CBX depletes multipolar and enriches for elongated GSCs. GSC morphotype distribution after 96h treatment with 125-500 µM TMZ and 60 µM CBX. N= 1 GSC line (P06); 3 biological replicates. Error bars, SEM. (J) VPF and CBX reduce GBO viability. Viability of GBOs metronomically treated with 60 µM CBX, 0.075 µM VPF, 500 µM TMZ and combination of all three drugs for 10 days. Viability measurements are normalized on DMSO control (100%). N= 5 patients (P23, P24, P25, P26, P27) in two independent experiments. Error bars, SEM. **(K-N)** VPF and CBX cancel the enrichment of chemoresistant morphotypes. (K-M) Representative microscopy images of GBOs (P24) treated as in (J) and stained for nestin (magenta) and SOX2 (yellow) to identify GSCs. Note that while in TMZ treated GBOs (L) GSCs appear longer and more branched than in DMSO (K), the GBOs treated with the combination of drugs (M) display numerous nonpolar GSCs. (N) Quantification of (K-M and S11H-I). Error bars, SEM. Two-way ANOVA with Sidak post-hoc test; ****, P<0,0001, N=5 patients (P23, P24, P25, P26, P27) in two independent experiments.

Gene set enrichment analysis (GSEA) further revealed a pronounced signature of brain tumor multidrug resistance (Keshelava et al., 2007) specifically in elongated and multipolar GSCs, but not in nonpolar cells (Fig. 5C). Although both elongated and multipolar GSCs shared some core resistance genes, we identified morphoclass-specific drivers of resistance (fig. S10A, B). Multipolar GSCs prominently expressed the mitochondrial channel VDAC1 (fig. S10A), previously linked to stemness and chemoresistance and representing a potential therapeutic target through metabolic reprogramming (Arif et al., 2017; Shteinfer-Kuzmine et al., 2017). Supporting this, we readily detected mitochondria within protrusions of multipolar GSCs (Fig. 5D and S10C), consistent with prior observations of mitochondrial exchange in connected GBM cells adapting to therapy (Pinto et al., 2021). Collectively, our data suggest that multipolar GSCs resist therapy through network-mediated mitochondrial interactions and calcium waves.

In elongated GSCs, the resistance signature featured the transcription factor TEAD1 (fig. S10B), implicated in the Hippo pathway and vessel co-option as a resistance mechanism (Barrette et al., 2022; Pichol-Thievend et al., 2024). We hence examined the expression of YAP, key Hippo mediator. Immunofluorescence analysis in our patient samples revealed marked YAP presence in elongated GSCs in both core and periphery GBM tissue (S7D). Specifically, peripheral vascular-associated elongated GSCs displayed active YAP expression in primary GBM samples (Fig. 5E), suggesting that elongated cells leverage vascular interactions not only for invasiveness (see Fig. 3J-L), but also to evade therapeutic targeting. Additionally, elongated cells specifically expressed resistance-related genes SOX4 and HIF1A (fig. S7B), known regulators of invasion, angiogenesis, and vascular remodeling (Yao et al., 2024; Zheng et al., 2021). These analyses allow us to hypothesize that elongated and multipolar GSCs utilize different chemoresistance mechanisms.

In light of the morphoclass-specific chemoresistance pathways, we next tested whether the YAP inhibitor Verteporfin (VPF) would sensitize elongated GSCs. By treating two patient-derived GSC lines with increasing concentrations of VPF we observed a drastic decrease in cell viability (Fig. 5F, fig. S11A, C), particularly driven by the specific depletion of elongated GSCs (Fig. 5G, fig. S11A, B, D, E). Moreover, the combination with TMZ, which preferentially targets simple morphoclasses, led to a further increase in cell death. To examine a potential synergistic effect between TMZ and VPF, we calculated the excess over Bliss score (Liu et al., 2018), which showed that the two drugs are highly synergistic (score of 8).

To target chemoresistant multipolar GSCs we sought to break network-mediated mitochondrial exchange and calcium waves as they likely mediate the chemoresistance of those cells (Pinto et al., 2021; Valdebenito et al., 2020; Weil et al., 2017). Considering that these processes are mediated by gap junctions (Hausmann et al., 2023; Pinto et al., 2021) (and see Fig. 4H), we treated two patient-derived GSC lines with the gap junction blocker CBX. We detected a striking synergistic effect of CBX with TMZ (excess over Bliss, score of 19) (Fig. 5H, S11F, G). At the highest concentration tested, CBX induced significant cell death, leading to a complete depletion of multipolar GSCs and selection of elongated GSCs, which constituted >80% of the remaining cells (Fig. 5H, I, S11F). This indicates that CBX treatment specifically targets multipolar GSCs.

To explore the effectiveness of these compounds in a physiologically more relevant model system, we treated patient-derived organoids with CBX, VPF, TMZ and the combination of the three. After 10 days of metronomic treatment, we observed a marked decrease in cell viability upon both VPF and TMZ treatments. Importantly, this effect was more pronounced upon the co-treatment with all the three drugs (Fig. 5J). In agreement with our data in GSC lines, CBX and VPF treatments reduced the proportion of multipolar and elongated GSCs, respectively (Fig. 5K-N, fig. S11H, I). Overall, our data suggests that GSCs have morphology-specific therapeutic vulnerabilities with nonpolar cells being sensitive to TMZ, elongated GSCs to YAP inhibitors and multipolar GSCs to gap junction blockers.

## Discussion

Our study establishes a direct link between GSC morphology, transcriptional identity and cellular behaviors, particularly invasiveness and chemoresistance, which are two key challenges in the clinical management of GBM. Moreover, we show that GSCs have morphoclass-specific therapeutic vulnerabilities opening the path to the development of cell morphology-informed therapies in GBM. While previous studies associated GBM cell morphology with proliferation, invasiveness and chemoresistance (Bhat et al., 2013; Monzo et al., 2016; Ratliff et al., 2023; Venkataramani et al., 2022b; Xie et al., 2014), we significantly extend this knowledge by specifically focusing on stem cell morphotypes and using 3D patient-derived organoids and assembloids. The integration of our custom spatial transcriptomics method, CellShape-seq, with functional assays and pharmacological manipulations, enabled us to map transcriptomic signatures onto individual GSC morphologies, providing a detailed understanding of how structural diversity is linked to cellular function and response to therapy.

We identified three distinct GSC morphoclasses - nonpolar, multipolar and elongated - that align with clinically relevant GBM subtypes defined by metabolic and developmental attributes (Bhat et al., 2013; Monzo et al., 2016; Ratliff et al., 2023; Venkataramani et al., 2022b; Xie et al., 2014). Nonpolar cells showed differentiation signatures, reduced proliferation and high sensitivity to TMZ treatment. In contrast, multipolar and elongated GSCs exhibited progenitor-like states, high proliferation and chemoresistance. Furthermore, we demonstrated that multipolar GSCs are interconnected in functional multicellular networks, while elongated GSCs primarily interact with blood vessels and are implicated in tumor invasiveness. Our findings suggest that morphological complexity, as observed in elongated and multipolar cells, is intrinsically connected to stemness and reliance on mitochondrial oxidative phosphorylation (OXPHOS). Given that nonpolar GSCs show differentiation signatures and are more sensitive to TMZ (Fig. 5A-C), it would be interesting to examine if OXPHOS inhibitors (Bernhard et al., 2023; Garofano et al., 2021) could potentially sensitize resistant elongated and multipolar GSCs by inducing their morphological shift towards the vulnerable nonpolar state.

Our data in tumor tissues and patient-derived organoids adds to our previous study (Barelli et al., 2025) showing that stem cell morphology is a new layer of GBM heterogeneity in clinically relevant models where the intricate interplay between GSCs and the microenvironment is largely recapitulated. GBM displays high inter- and intra-tumoral heterogeneity at the genomic, transcriptomic, epigenetic and metabolic levels (Barelli et al., 2025; Osswald et al., 2015; Weil et al., 2017), but the cell biological relevance of this remains largely unexplored. Here, we have linked specific cell morphologies to different transcriptional identities and cell states and showed that GSC morphoclasses predict specific oncological functions. An important question emerging from our data is whether morphological states are stable or flexible throughout tumor evolution and therapy exposure. Our previous findings indicate stable inheritance of GSC morphology (Barelli et al., 2025), yet environmental conditions can also induce morphological plasticity (Xie et al., 2014). Future research should explore how therapeutic interventions might drive morphological transitions and whether leveraging these shifts could enhance treatment responses.

To connect individual cell morphology with transcriptional states, we developed CellShape-seq, a custom spatial transcriptomics platform enabling morphotype-specific transcriptomic profiling. Current spatial biology methods primarily provide tissue-level spatial resolution, while single-cell approaches lack direct morphological integration (Longo et al., 2021; Moses and Pachter, 2022; Williams et al., 2022). CellShape-seq fills this gap by directly linking cell shape and transcriptional identity (Fig. 2A), thus allowing us to map cell states and pathways onto specific stem cell morphotypes. This technology offers applications beyond GBM. For example, breast cancer cells are known to exist in different morphologies associated to different clinical outcomes (Alizadeh et al., 2020; Conner et al., 2024; Sali et al., 2024), but their morphotype-specific transcriptome has not been characterized. In neuroscience, the advent of patch-seq has allowed to transcriptionally map neurons based on their morphology, localization and electrophysiology but it has a low throughput and has limited application to other cell types (Cadwell et al., 2016). CellShape-seq can hence be useful across biomedical disciplines to disentangle the relationship between cell morphology, transcriptional signatures and cell function.

From a therapeutic standpoint, our findings highlight the necessity of integrating morphological analyses into clinical diagnostics and personalized treatments. Importantly, our data indicate that the chemoresistance mechanisms differ notably between morphoclasses, which underscores the need for combinatorial treatments simultaneously targeting different aspects of the tumor (Ghosh et al., 2018). Multipolar GSCs formed interconnected networks mediated by calcium signaling, gap junctions, and likely mitochondrial exchange via cell protrusions, consistent with previous reports (Barelli et al., 2025; Osswald et al., 2015; Weil et al., 2017). Such networks are a paradigmatic example of how a morphological feature – the presence of cellular protrusions and thus intercellular connections – directly augments GSC survival as cells collectively resist therapies that would kill them individually. In line with current clinical trials repurposing anti-epileptic agents (Gonzales et al., 2024) and gap junction blockers (Schmidt et al., 2025) for GBM, we show that blocking gap junctions disrupts such networks and sensitizes multipolar GSCs to standard chemotherapeutics. Furthermore, we identified that these cells express high levels of the mitochondrial channel VDAC1, a promising target for metabolic reprogramming (Arif et al., 2017; Shteinfer-Kuzmine et al., 2017). Meanwhile, the chemoresistance signature of elongated GSCs features an activation of the YAP/TAZ/TEAD pathway, which has been previously linked to GBM invasiveness and vascular co-option (Barrette et al., 2022; Pichol-Thievend et al., 2024). We showed that inhibitors such as Verteporfin, targeting YAP signalling, sensitise elongated GSCs but it remains to be determined whether they would concurrently also impair invasiveness in these aggressive cells. Besides, as interaction with the vasculature seems crucial to mediate both invasiveness and chemoresistance, pharmacological agents that disentangle GSC-vasculature crosstalk would also hold potential to reduce both.

In summary, our findings demonstrate that the intrinsic variability in GSC morphology is linked with critical aspects of GBM aggressiveness and with different windows of therapeutic vulnerabilities. This shows the necessity of moving beyond traditional cytotoxic treatments towards combinatorial treatment strategies that target the physical and phenotypic adaptations of the tumor. Morphology-informed therapies could show promise in this context. We hence propose that integrating these morphological insights into treatment design might improve responses to therapy and patient outcomes.

## Methods

### Human samples

Glioblastoma patient samples were obtained from Ospedale Nuovo di Legnano following informed patient consent. A total of 21 patient samples from both males and females between 37 and 87 years old with a diagnosis of grade IV astrocytoma were included (see Table S2). Fresh surgical resections were collected in Hibernate™-A Medium (A1247501, Thermo Fisher) containing Penicillin/Streptomycin 100X (ECB3001D, Euroclone) and Amphotericin B 100X (15290018, Thermo Fisher) and transported to Human Technopole at 4°C for processing. Immediately upon arrival, the tissue was dissected in Hibernate™-A Medium and washed with the same medium to remove cellular debris. For each patient sample most of the tissue was further processed to generate patient-derived glioblastoma organoids, some was snap frozen and the remaining was fixed in 4% paraformaldehyde (PFA) After 24h of fixation, the tissue was left in 15% and 30% sucrose gradients for 24 h each. Following embedding in OCT compound Killik (05-9801, Bio Optica), serial sections of 20 µm were cut at the cryostat and stored at −20°C for immunofluorescence experiments. For most patients we obtained both core and peripheral tissue, with the core exhibiting high and the periphery low 5-ALA signal intensity during surgical resection. From those tumor samples we generated both organoids and fixed frozen sections for immunofluorescence.

### GBM organoid generation and culture

The dissected surgical specimens were cut into small pieces (0.5 to 1 mm diameter) with microdissection scissors and incubated in 1X RBC Lysis Buffer (00-4333-57, Thermo Fisher) for 10 minutes in shaking, following the previously established protocol (26). After washing with HibernateTM-A medium, the dissected pieces were cultured in 6-well low attachment plates (Corning, 3471) in their GBO medium composed of 50% DMEM:F12 (Thermo Fisher Scientific, 11330057), 50% Neurobasal (Thermo Fisher Scientific, 21103049), 1X GlutaMax (Thermo Fisher Scientific, 35050038), 1X NEAAs (Thermo Fisher Scientific, 11140050), 1X PenStrep (ECB3001D, Euroclone), 1X N2 supplement (Life Technologies, 17502-048), 1X B27 (Thermo Fisher Scientific, 17504001), 1X 2-mercaptoethanol (Sigma Aldrich, M6250-100ML), and 2.5 μg/ml human insulin (Sigma Aldrich, I0516) and placed on an orbital shaker rotating at 120 rpm within a 37°C, 5% CO2, and 90% humidity sterile incubator. The organoids generally form within 2-4 weeks and were then used for experiments up to week 15 of culture.

### GBM stem cell culture

Onda 11 cells were reconditioned to glioblastoma stem cells (Onda 11 GSCs) and grown on Laminin (5ug/ml, Sigma, L2020)-coated plates in serum-free media (GSC medium) composed of DMEM/F-12 with 15 mM HEPES and L-Glutamine (Thermo Fisher, 11330057), P/S, N2 supplement (Thermo Fisher, 17502-048), B27 supplement (Thermo Fisher, 17504-044), EGF (10µg/µl) and FGF2 (10µg/µl), as previously described (Barelli et al., 2025).

Primary patient-derived GSCs were derived from GBM organoids by dissociating them with Animal Free Collagenase/ Dispase Blend II (Sigma, SCR140) 50% diluted with GSC media. After cutting GBOs in small pieces, they were left in the dissociation solution at 37C for 30 min-1 h with 120 rpm shaking until most of the tissue was dissociated into single cells. Following washing, cells were plated on Laminin coated plates in GSC media as described above.

### Immunofluorescence

For immunofluorescence of human glioblastoma tissue and organoids, antigen retrieval was performed by incubating the slides for 45 min with 10 mM citrate buffer pH6.0 in 70°C oven. Following three washes in PBS, the tissue was permeabilized for 30 min in 0.3% Triton X-100 at room temperature, quenching for 30 min in 0.1 M glycine in PBS at room temperature and blocking for 30 min in blocking solution containing 10% normal donkey serum (Jackson ImmunoResearch, 017-000-121), 300 mM NaCl, 0.5% Triton X-100 in PBS. Primary antibodies (see Table 1) were incubated in blocking solution for 12 h, at 4°C. Following three washes in PBS, the sections were incubated with secondary antibodies Alexa Fluor PLUS (1:500, Thermo Fisher) and DAPI (1:2000, Thermo Fisher, 62248) in 0.3% Triton X-100 in PBS for 1 h at room temperature an temperature and washed again 3 times in PBS and mounted on microscopy slides with Mowiol 4-88 (Sigma Aldrich, 81381) + DABCO (antifade, Sigma Aldrich, D27802).

### Microscopy and image analysis

Microscopy of fixed samples was performed a Zeiss Axioscan z.1 slide scanner for overview images of whole organoids and confocal microscopes Zeiss LSM980 point-scanning confocal or LSM980-NLO point-scanning confocal or a Nikon Ti2-CREST spinning disk confocal microscope for other images. The confocal images were acquired with a PlanApo 10X/0.45 dry or a PlanApo 20X/0.8 dry or a PlanApo 40X/1.4 NA oil immersion objectives using 405nm, 488 nm, 561 nm, 639 nm laser lines. The software used for all acquisitions was Zen Blue 3.7 (Zeiss) or NIS-Elements (Nikon). Once the parameters of acquisition for control conditions had been defined, they were kept constant for all the samples within the same experiment.

For live imaging and the associated analysis see below “Calcium imaging and analysis”. All manual cell quantifications were performed in Fiji ImageJ using the CellCounter function, processed with Microsoft Excel, and plotted in GraphPad Prism. Morphotypes and morphoclasses were assigned for GSCs identified as Nestin+ SOX2+ cells in both primary tissue and organoids. For quantification of GSCs associated to blood vessels, we considered all Nes+ SOX2+ cells whose Nes+ was in contact with Laminin+ vessels. For assembloid invasion, we assigned morphoclasses to all EdU+ Nes+ SOX2+ cells in the core of the assembloid and in the invaded periphery. For histological sections, the labelling of core and peripheral tumor tissue came from the surgeons and the pathologists. For quantification of resistant Onda-11 GSC populations, morphoclasses were assigned to the cells at day 7 using bright field images and following the morphological classification we have established previously (16). For the TMZ experiments in organoids, we quantified the morphoclass distribution of Nes+ SOX2+ cells after 10 days of metronomic TMZ administration.

### CellShape-seq customized GeoMX spatial transcriptomics

Four GBM organoids from each of the four different selected patients were formalin fixed paraffin embedded (FFPE). After fixation in 10% NBF (Bio-Optica, 05-01004F) for 24 hours at room temperature and washes in PBS, the organoids were embedded in 2% low melting agarose (Invitrogen, 16520050). For paraffin embedding, the automatic tissue processor Sakura Tissue-Tek VIP6 was used. The paraffin blocks were cut on the microtome at 4 µm thickness. Slide preparation for GeoMX was carried out as instructed by the GeoMx DSP Manual Slide Preparation (MAN-10150-06). Briefly, the FFPE organoid sections were deparaffinized and rehydrated with Xylene, 100% EtOH, 95% EtOH, and DEPC-treated water. Antigen retrieval was performed with Tris-EDTA pH9 for 10 minutes at 99°C, followed by the exposure of RNA targets through Proteinase K 0,1 µg/ml for 15 min at 37°C. The sections were post-fixed in 10% NBF (Bio-Optica, 05-01004F) before the in situ hybridization with the Human Whole Transcriptome Atlas probe-set (GMX-RNA-NGS-HuWTA-4) would take place overnight in the hybridization oven (HybEZ II, ACDbio). Stringent washes were performed in a solution of 2X SSC and 50% formamide to remove off target probes. After blocking the non-specific staining with Buffer W provided by the company, the morphology markers were added by diluting the primary antibodies Anti-Nestin (1:500, Invitrogen, MA1-110) and Anti-SOX2 (1:200, R&D AF2018) in the same buffer. Following 1-hour incubation at room temperature the sections were washed in 2X SSC and the secondary antibodies Alexa Fluor PLUS Anti-Goat 555 (1:500, Invitrogen, A32816) and Anti-Mouse 647 (1:500, Invitrogen, A32787) along with the nuclear marker SYTO13 were added for 30 minutes at room temperature. Following washes in 2X SSC, the slides were stored in 2X SSC overnight at 4°C.

The following morning the slides were loaded and scanned by the NanoString GeoMX DSP machine. 16 ROIs (regions of interest) of the maximum size 660×785 µm (one per each organoid covering almost the entire organoid) were drawn, downloaded and opened into FiJi. After adjusting the scale to the same as the GeoMX platform, the AOI (area of illumination) single-cell segmentations were drawn using the free hand line tool around each of the desired Nestin+ SOX2+ cell. For each ROI, three separate masks, each for one of three morphoclasses (nonpolar, elongated, multipolar) was created, saved as .jpg files and re-uploaded onto the GeoMX platform. This resulted in 16 ROIs each one with three different AOI segmentations corresponding to the three morphoclasses of interest within each organoid.

### Library preparation and sequencing

Oligos from selected AOIs were collected into a 96 well plate and were amplified and tagged with ROI-specific barcodes during library preparation, adding sequencing adapters for subsequent high-throughput sequencing, following the protocol described in the GeoMX DSP NGS Readout User Manual (MAN-10153-06). Libraries were sequenced on a NextSeq2000 instrument (Illumina) generating per each AOI a number of reads pairs derived by applying the following formula n. read pairs = total collection area (µm2) * sequencing depth factor (equal to 100 for the WTA panel).

### Spatial transcriptomics sequencing analysis

Raw sequencing reads were processed using the GeoMx NGS Pipeline (NanoString, version 2.3.3.10) for alignment, deduplication, and quantification of the probes from the GeoMx Human Whole Transcriptome Atlas. To account for differences in sequencing depth, count data were normalized across AOIs using size factors computed via the trimmed mean of M values (TMM) method (71). Principal Component Analysis (PCA) was performed on the standardized data using scikit-learn v1.3.2 [https://scikit-learn.org/].

Differential expression analysis was conducted using PyDESeq2 v0.4.1 (72) on raw counts. A model was fitted, incorporating the patient of origin of the organoid and the AOI morphoclass as covariates. Gene expression contrasts were computed for each morphoclass (nonpolar, elongated, and multipolar) against the other two. Genes with an adjusted p-value < 0.05 were considered significantly differentially expressed. Significantly differentially expressed genes were functionally characterized through over-representation analysis of Biological Process Gene Ontology (GO) terms (2023 version (73)). Enrichment analysis was performed using the enrichr function from gseapy (74) to perform Over-Representation Analysis (ORA) and Gene-set Enrichment Anlaysis (GSEA), respectively.

Functional analysis of the DEGs was performed using QIAGEN Ingenuity Pathway Analysis (IPA) (75). Gene expression data from elongated and multipolar cell lines (gene identifiers and corresponding measurement values) were uploaded into the application. Each identifier was mapped to its corresponding entity in QIAGEN’s Knowledge Base. Molecules whose expression was significantly perturbed, called Network Eligible molecules, were overlaid onto a global molecular network developed from information contained in the QIAGEN Knowledge Base. Networks of Network Eligible Molecules were then algorithmically generated based on their connectivity.

### Biological pathway deconvolution analysis and GBM subtypes alignment

To further characterize the transcriptomic dataset, a pathway deconvolution analysis was performed following an approach similar to that described in Garofano et al. (9). The main points of this approach are:

1. Reduce inter-patient variability by standardizing gene expression within each patient group.
2. Perform single-sample pathway enrichment analysis using the Mann–Whitney– Wilcoxon Gene Set Test (MWW-GST).

More specifically, for each AOI gene expression was standardized within the patient subgroup to generate an AOI-specific ranked list of genes. A collection of gene sets was curated by merging: Hallmark, Gene Ontology, Oncogenic Signatures, KEGG MEDICUS, and Reactome pathways from the Molecular Signatures Database (MSigDB) (76). Gene sets containing fewer than 6 or more than 200 genes were excluded.

For each AOI, the Normalized Enrichment Scores (NES) for all selected gene sets were computed using a Mann-Whitney U test, comparing the ranks of genes in each pathway to background genes. The NES represents the probability that, in a given AOI, a randomly chosen gene from the gene set ranks higher than a randomly chosen background gene. The NES was calculated from Mann-Whitney U using the following formula: NES = U/nm with n being the number of genes in the gene list and m being the number of background genes. With each AOI now represented by a vector of NES values, a pathway-based embedding of the dataset was generated through dimensionality reduction. The 500 gene sets with the highest dispersion in NES values were selected. In this reduced space, the dataset was represented first by a nearest-neighbors graph constructed using k-nearest neighbors (k-NN) and then by a Uniform Manifold Approximation and Projection (UMAP) embedding, both obtained using functions from scanpy (77). Clustering was then performed using the Leiden algorithm with a standard resolution of 1.0, grouping AOIs based on NES similarity.

To further characterize the identified clusters in the context of glioblastoma transcriptomic profiles, we generated functional scores based on the classification proposed in (9). Specifically, we extracted four sets of gene pathways corresponding to the biological programs described in their single-cell analysis of glioblastoma tissue: Proliferative/Progenitor, Neuronal, Mitochondrial, and Glycolytic/Plurimetabolic. For each AOI, we computed a functional score for each of these biological classes by averaging the NES of the associated pathways. The final scores were computed using the score_genes function from scanpy.

### Assembloid invasion assay

Assembloids were generated by fusing a core and a peripheral GBM organoids from the same patient. The two organoids were kept in a 1.5 ml Eppendorf for 5-7 days until when they were completely fused, meaning that they would not break when pipetting up and down the media. After fusion the assembloids were transferred into a low-attachment 6-well or 24-well plate until fixation in 4% PFA.

For invasion assays, before assembloid formation the core GBM organoids were treated with EdU for 24 hours so that multiple cells would uptake EdU. Following washes in PBS, they were fused with their corresponding peripheral organoid as described above. At day 15 (5 days in Eppendorf) + 10 days in low attachment plates, i.e. 10 days post fusion), the assembloids were fixed and processed into 30µm cryosections as described above to analyse the EdU+ core cells which had invaded the peripheral part of the assembloid. For EdU detection by immunofluorescence, Click-iT™ EdU Cell Proliferation Kit for Imaging, Alexa Fluor™ 647 dye (Thermo Fisher, C10340) was used. These invasion assays were performed on two different assembloid batches, each containing at least four patients with 2-4 assembloids per patient.

### Calcium imaging and analysis

GBM organoids were treated with Fluo4 AM Ca2+ indicator (Invitrogen, F14201) at 2 µM diluted in GBO media for 2 hours. Following three washes in GBM organoid media, the organoids were transferred to an IBIDI µ-Slide 8 Well and taken to the Nikon Ti2 - CrestOptics V3 spinning disk microscope for live imaging. To attach the organoids to the bottom of the dish, a homemade manufactured anchor was put on top of the organoid for the duration of the movie. Live imaging was performed at 37 °C with 5% CO2 and 3-5 minutes widefield movies were taken with a PlanApo 20X/0.8 dry using 488 nm laser lines with an acquisition frame rate of 5.05 frames/second. 2-6 organoids for each of the six different patients were used to obtain 1-4 movies per organoid.

To inhibit the calcium activity, a gap junction blocker Carbenoxolone (CBX) (Sigma Aldrich, C4790) was incubated at 100 µM for 15 minutes, after which another 3-5 minutes movie was taken in the same field of view where the activity was originally recorded.

The script for the analysis of calcium activity was adapted from (48). Briefly, ROIs of single cells were automatically segmented using a Fiji macro with the following steps “Auto Local Threshold-Otsu method, radius 35”> “Watershed” > “Analyze Particles-size 60-500”. Each movie was bleach-corrected for exponential decay, and the mean intensity of each ROI was computed over all time frames (Fiji). After assigning each cell to its morphoclass, we proceeded with the network analysis in MATLAB (MathWorks Inc). The calcium transient detection method was modified from the original script: the fluorescent values were smoothed using a moving average over 7 points, and the baseline fluorescence (F0) was calculated by identifying the minimum value within a 100-points window preceding the time point of interest. Finally, ⍰F/F0 (DFF) was calculated. A spike was detected when the peak height exceeded 4 times the standard deviation of all DFF values in the movies. The parameters of the network analysis were adapted from (48) as follows: cells were considered active with 1 peak within our 3 minutes movies; periodic cells were defined with more than 3 firings within the 3 minutes movies; trigger cells must excite at minimum 2 cells and hub cells must correlate with more than 4 cells.

### Pharmacological treatments and resistance assays

Onda-11 GSCs were metronomically treated with 200 µM Temozolomide (TMZ) (Sigma Aldrich, T2577) every 48h for 7 days to select the resistant GSC morphoclasses. Bright field images were taken to quantify the proportion of GSC morphoclasses at day 7. Primary GSCs were treated with 0.025-0.1 µM Verteporfin (VPF, Sigma, SML0534), 20-60 µM Carbenoxolone Disodium Salt (CBX, Sigma, C4790-5G) alone or in combination with 125-500 µM TMZ for 96h, when brightfield images were taken to analyze their morphologies. For all 2D cell lines, the brightfield images were taken at EVOS Cell Imaging Systems (Thermo Fisher), cell viability was measured with the RealTime Glo MT Viability Assay (Promega, G9711) and luminescence was detected with GloMAX Discover (Promega).

GBM organoids were treated with TMZ for 96 h with doses ranging from 200 µM to 5000 µM to test the most effective doses. To select the GSC population resistant to TMZ, GBM organoids were metronomically treated with 200 µM, 500 µM Temozolomide (TMZ) (Sigma Aldrich, T2577) or DMSO in GBM organoid medium every 48 hours for 10 days. To examine their effects on GBM organoids, 0.075 µM VPF and 60 µM CBX were metronomically administered either alone or in combination with 500 µM TMZ. After 10 days of all treatments, the GBM organoids were fixed in 4% PFA and processed into 20 µm cryosections for immunofluorescence and GSC morphological analysis. GBM organoid viability was assessed with the CellTiter-Glo® 3D (Promega, G9681) and the luminescence was normalized on the area of the organoids and captured with GloMAX Discover (Promega).

### Statistical analysis

All statistical analyses were conducted using Prism (GraphPad Software) or Microsoft Excel. To test for statistical significance (p<0.05), one-way or two-way ANOVA with Sidak or Tukey post hoc tests, Fisher exact test and Student’s t test were used. For each graph, the number of samples and patient codes, error bars, statistical test and the p-value are noted in the figure legends.

To calculate the combinatorial effect of pharmacological compounds, we used the “Excess over Bliss” (EoB) score which quantifies the difference between the observed effect (E_observed_) and the expected additive effect (E_expected (Bliss)_), where the E_expected (Bliss)_ is:

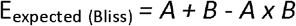

When the EoB score is null *A* and *B* have an additive effect, when EoB > 0 there is synergism between the two compounds and when EoB < 0 *A* and *B* are antagonists.

## Acknowledgments

We are grateful to the services and facilities of HT for the outstanding support provided, notably, N. Maghelli and the team of the National Facility (NF) for Light Imaging, including the infrastructural units (IU) for Ion imaging and Tissue processing, A. Riva and the team of the NF for Data Handling and Analysis, including D. dalle Nogare and the team of the IU for Bioimage analysis, and the team of the NF for Genomics. We thank all members of the Kalebic laboratory for their support and discussions. CB is a Ph.D. student within the European School of Molecular Medicine (SEMM).

## Author contributions

Conceptualization: CB, NK

Methodology: CB, MB, FM, DR, IB, CP, SF

Investigation: CB, FM, NA, VS, SF

Formal analysis: CB, MB, FM, RB

Visualization: CB, MB, FM, RB

Resources: AC, GMS, RS

Funding acquisition: NK

Project administration: NK

Supervision: NK

Writing – original draft: CB, NK

Writing – review & editing: all authors

## Data and materials availability

The sequence data has been deposited at SRA with the accession number: BioProject ID PRJNA1251971.

## Supplementary Materials

Figs. S1 to S10

Tables S1 to S2

Movies S1 to S4

