## Supplemental figures and legends for "Stem cell morphology defines functional heterogeneity and therapeutic vulnerabilities in glioblastoma"

Supplemental information

**Supplementary figures**

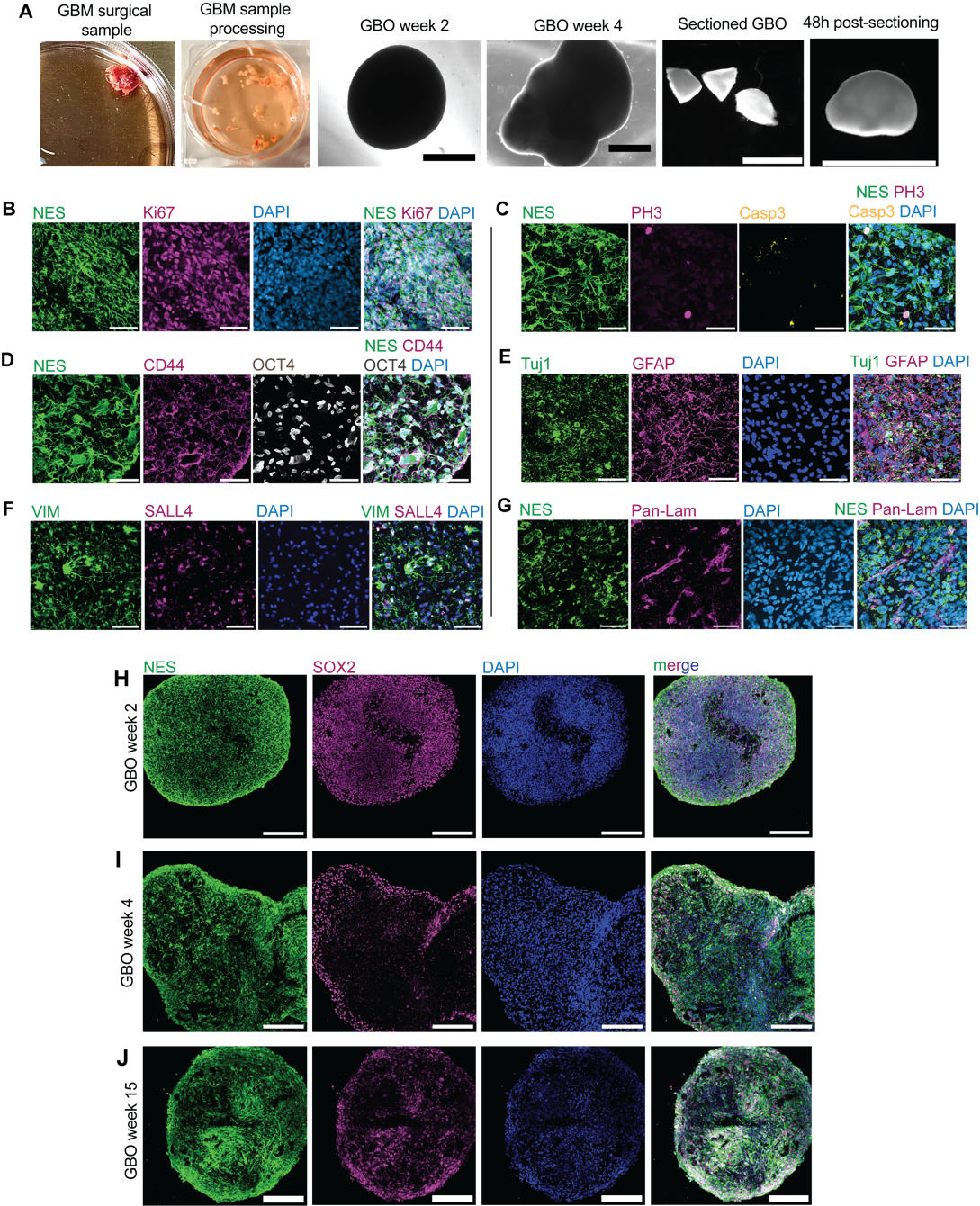

### **Figure S1. GBM organoids (GBO) culture and characterisation.**

**(A)** Schematics of GBO formation and culture starting from GBM surgical sample to GBO formation, growth, sectioning and regrowth. Photographs (first two) and bright-field images (remaining four). Scale bars, 500 µm. **(B-G)** GBO characterisation for various GBM and GSC markers. GBOs express the GSC markers Nestin (B, C, D, G), CD44 (D), OCT4 (D), SALL4 (F), (see also SOX2 in Fig.1); the GBM cell marker and neuronal marker TuJ1(E), the GBM cell and astrocyte markers GFAP (E) and Vimentin (F); the mitotic marker PH3 (C) and marker of cycling cells Ki67 (B); the blood vessel marker Pan-Laminin (G); and have low expression of the apoptotic marker Cleaved-Caspase 3 (C). Scale bars, 100 µm. (**H-J**) Representative microscopy images of GBOs at culture week 2 (H), 4 (I) and 15 (J), expressing the GSC markers Nestin and SOX2 and displaying GSC morphological heterogeneity (see Fig. 1). Scale bars, 500 µm.

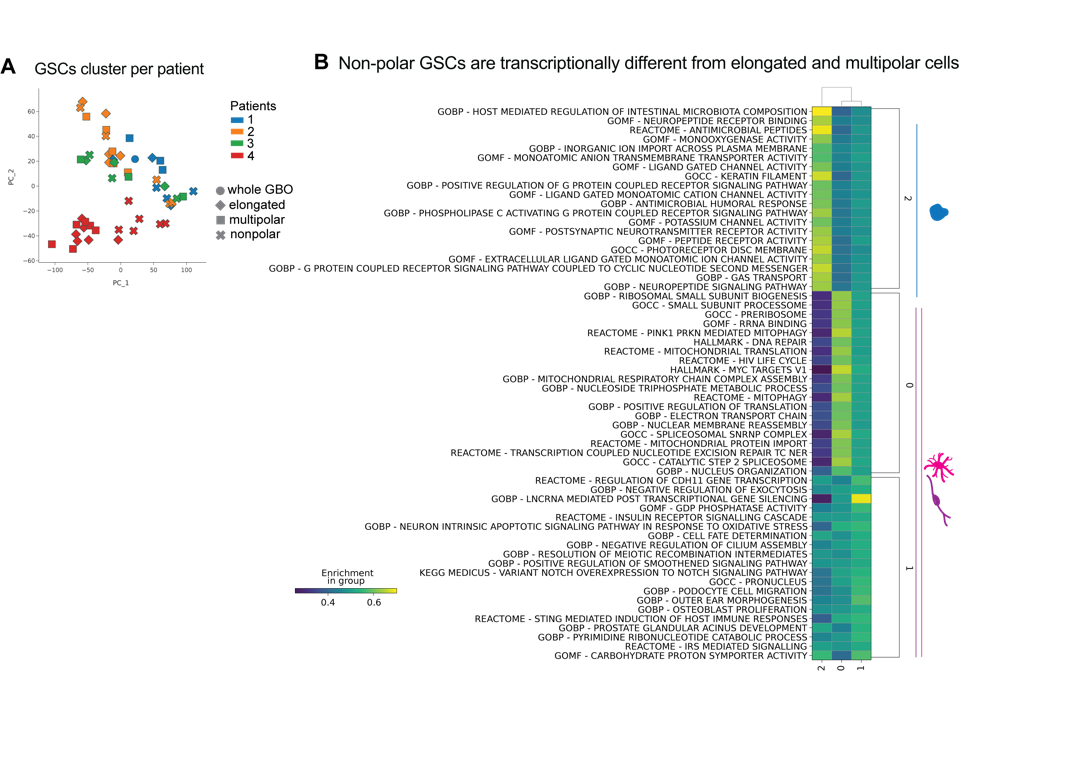

### **Figure S2. GSCs have morphoclass-specific transcriptomic signatures.**

**(A)** Dimensionality reduction of the GeoMX data. Principal Component Analysis (PCA) plot of the expression profiles of the AOIs from the GeoMX experiment on the GBM organoids. Colors indicate the patient from whom the organoid was derived, while symbols represent different morphoclasses (gray in the legend). **(B)** Pathway deconvolution cluster unbiased characterization. Enrichment values for the 20 most highly enriched pathways in each Leiden cluster compared to the rest of the dataset. For each cluster, the top pathways were selected based on a ranking determined by a Wilcoxon rank-sum test (implemented in the *rank_genes_groups* function of scanpy).

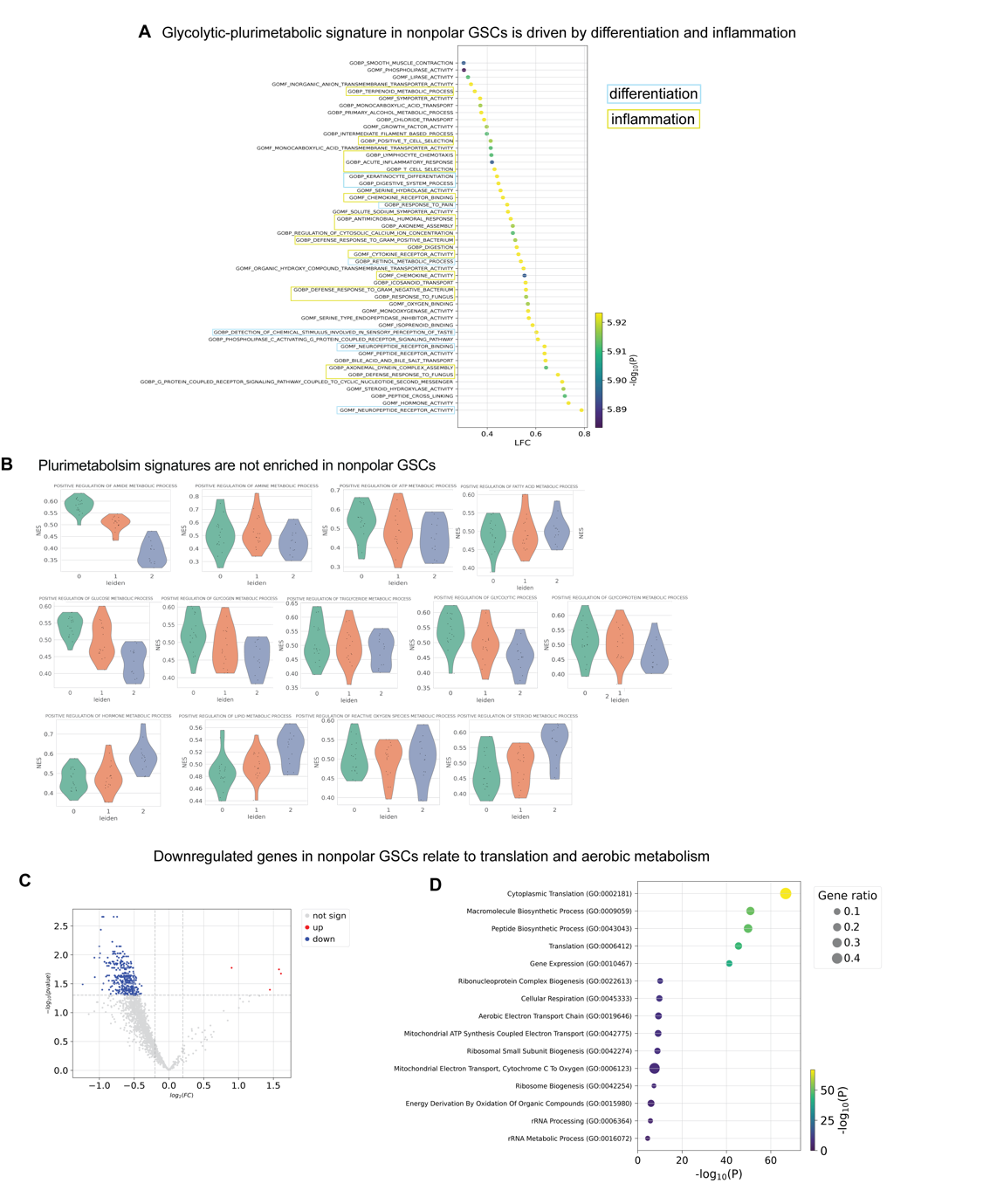

### **Figure S3. Nonpolar GSCs have transcriptomic signatures of differentiation, inflammation and hormonal metabolism.**

**(A)** Enrichment values for the 50 most highly enriched pathways for the Glycolytic-Plurimetabolic signature in Leiden cluster 2 (comprised of 80% nonpolar GSCs, see Fig. 2E). The top pathways were selected based on a ranking determined by a Wilcoxon rank-sum test (implemented in the *rank_genes_groups* function of scanpy). Note that most terms relate to differentiation (highlighted in blue) and inflammatory pathways (yellow). **(B)** Functional scores of the pathway deconvolution clusters. Violin plots depicting different functional scores of pathways related to plurimetabolism (amide, amine, ATP, carbohydrate, cholesterol, collagen, fatty acid, glucose, glycogen, glycoprotein, hormone, lipid, reactive oxygen species, steroid and triglyceride metabolic process) grouped by Leiden clusters. Functional scores for each AOI were computed by averaging pathway enrichment values over the pathway classes described in ^1^. **(C)** Differential expression analysis (DEA) of nonpolar cells vs the rest. Volcano plot showing the results of a DEA, comparing the expression profiles of AOIs with non-polar cells against the rest of the dataset. Differential expression analysis was performed using PyDESeq2. The x-axis represents Log-Fold Change (LFC), while the y-axis shows the negative log10 of the adjusted p-value. Dots highlighted in red are significantly up-regulated genes (LFC > 0 and adjusted p-value < 0.05), while dots highlighted in blue are significantly down-regulated genes (LFC < 0 and adjusted p-value < 0.05). Top upregulated genes: *FUCA1*, *HSPA1A*, *GPNMB*. **(D)** Top 15 downregulated pathways in nonpolar cells. Significantly downregulated pathways identified through Over-Representation Analysis (ORA) using the *enrichr* function from *gseapy*. The analysis was performed on genes with significantly lower expression in ROIs containing non-polar cells compared to the rest, selected based on a Log-Fold Change (LFC) < 0 and an adjusted p-value (corrected for multiple testing) < 0.05. In the plot, the x-axis position and dot color indicate the significance level of pathway enrichment, while dot size represents the proportion of differentially expressed genes associated with each GO term.

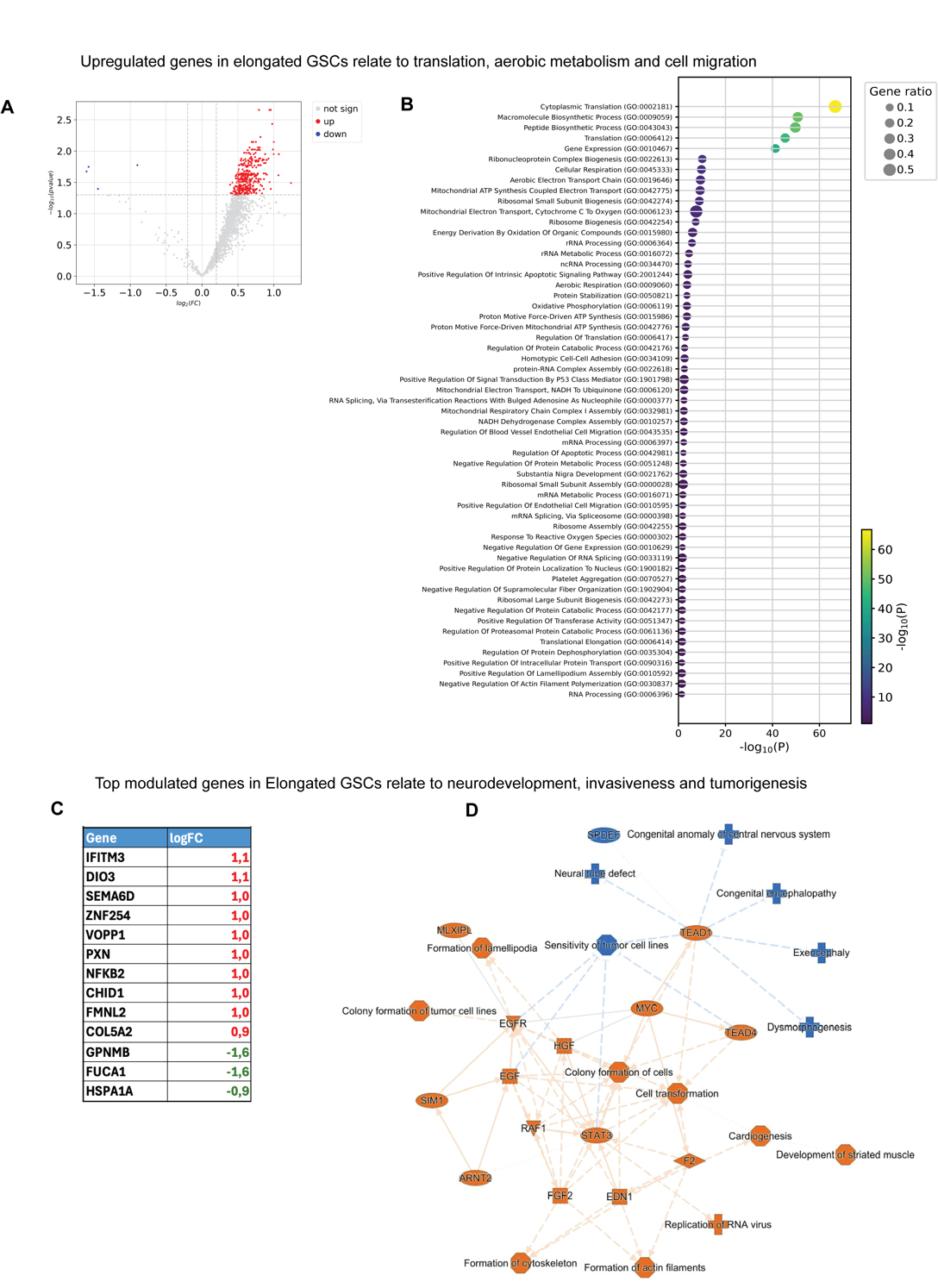

### **Figure S4. Elongated GSCs are enriched in pathways related to neurodevelopment and cell migration.**

**(A)** Differential expression analysis (DEA) of elongated cells vs the rest. Volcano plot showing the results of a DEA, comparing the expression profiles of AOIs with elongated cells against the rest of the dataset. Differential expression analysis was performed using PyDESeq2. The x-axis represents Log-Fold Change (LFC), while the y-axis shows the negative log10 of the adjusted p-value. Dots highlighted in red are significantly up-regulated genes (LFC > 0 and adjusted p-value < 0.05), while dots highlighted in blue are significantly down-regulated genes (LFC < 0 and adjusted p-value < 0.05). **(B)** Upregulated pathways in elongated cells. Significantly upregulated pathways identified through Over-Representation Analysis (ORA) using the *enrichr* function from *gseapy*. The analysis was performed on genes with significantly higher expression in ROIs containing elongated cells compared to the rest, selected based on a Log-Fold Change (LFC) > 0 and an adjusted p-value (corrected for multiple testing) < 0.05. In the plot, the x-axis position and dot color indicate the significance level of pathway enrichment, while dot size represents the proportion of differentially expressed genes associated with each GO term. See also Fig. 3A. **(C)** List of the top modulated genes with corresponding log fold change values observed in Ingenuity Pathway Analysis (IPA) Core Analysis for elongated cells. Note that out of 10 top genes, seven are associated to neurodevelopment, axon growth and tumor cell migration (SEMA6D, DIO3, VOPP1, PXN, CHID1, FMNL2 and COL5A2) **(D)** Graphical summary providing an overview of the major biological themes observed in IPA Core Analysis for elongated cells and their interrelations. It highlights the most significantly activated (orange) or inhibited (blue) canonical pathways (hourglass), diseases (cross), biological functions (octagon), and upstream regulators (rhombus for enzymes, square for cytokines, hexagon for translational regulators, oval for transcriptional regulators, inverted triangle for kinases, circle for others). The summary is constructed using machine learning techniques to prioritize and connect entities. Direct interactions are shown with solid lines, indirect interactions with dashed lines, and inferred edges with dotted lines.

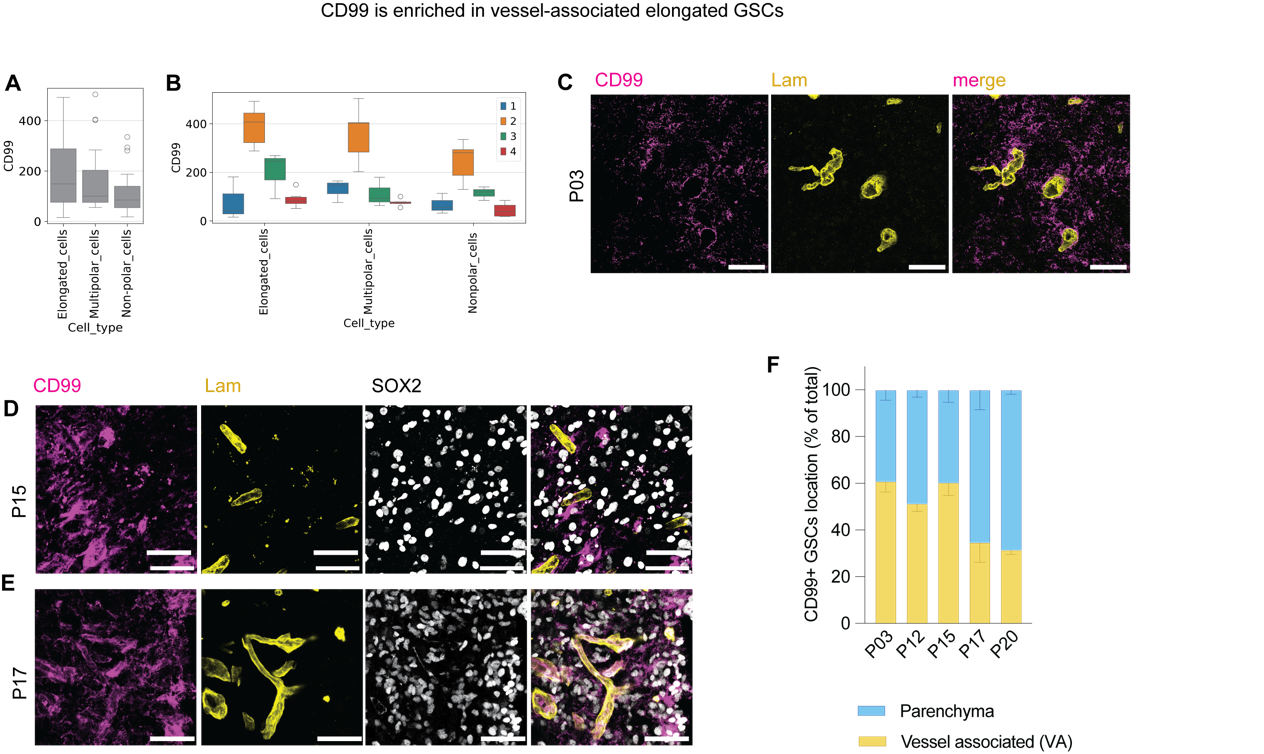

### **Figure S5. CD99 is a putative marker of elongated GSCs and it is enriched in vessel-associated GSCs.**

**(A, B)** CD99 is a putative marker of elongated GSCs. Expression level of CD99 across cell types (A) and patients (B). Box-and-whisker plots showing the normalized expression levels of CD99 from the GeoMX AOIs across cell type (A) and cell type and patient (B). The box shows the dataset quartiles, while the whiskers extend to the rest of the distribution, except for “outlier” points that are below/above the first/third quartile with a distance of more than 1.5 times the interquartile range. **(C-E)** Representative microscopy images of patient GBM tissue stained with CD99 (magenta), SOX2 (white in D and E) and Laminin (yellow) from 3 different patients (P03, P15, P17). Scale bars, 100 µm. **(F)** Quantitative distribution of CD99+ GSCs associated to blood vessels (yellow) or in the parenchyma (blue). Error bars, SEM. N=5 patients (P03, P12, P15, P17, P20).

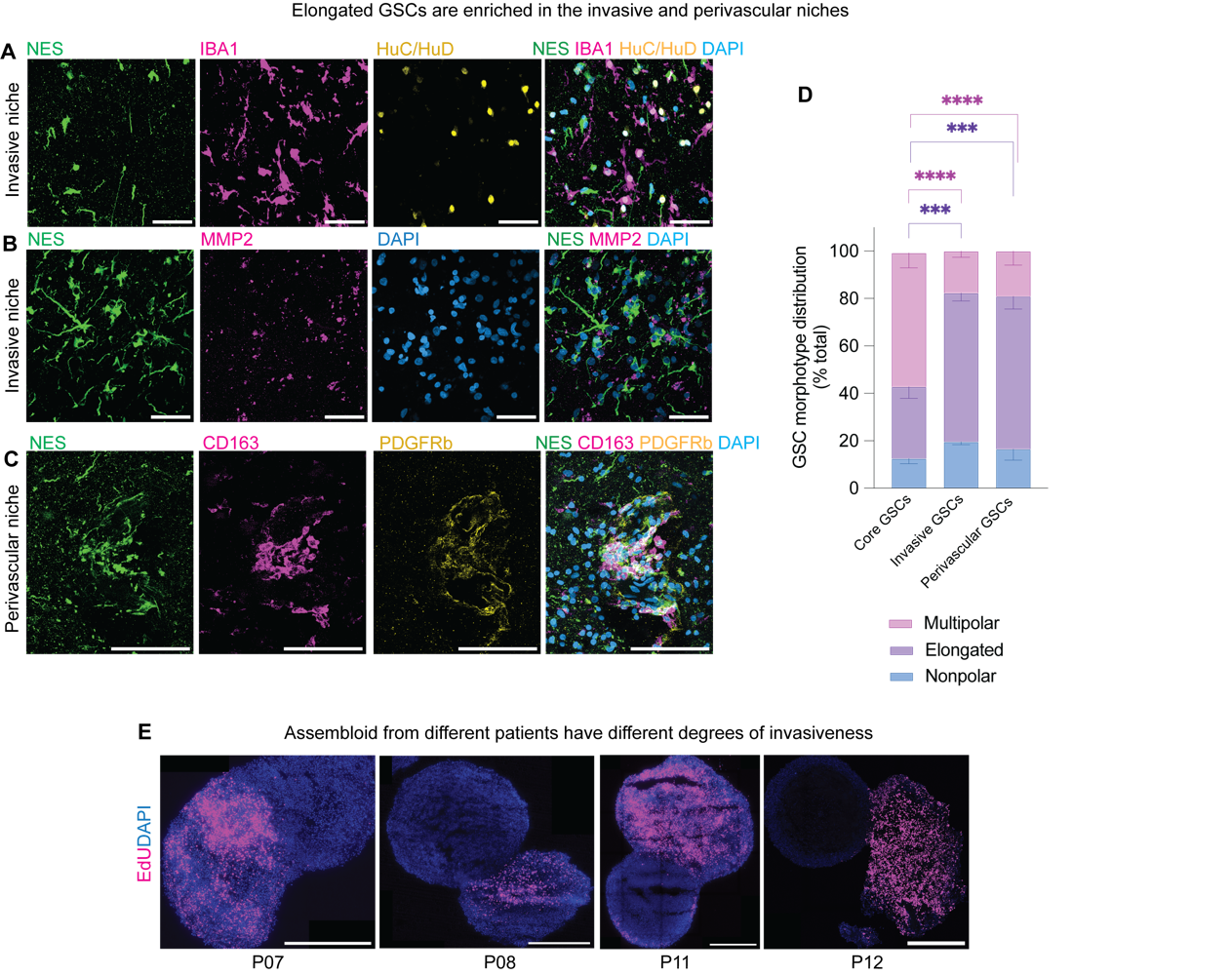

### **Figure S6. Elongated GSCs are invasive and perivascular.**

**(A-D)** Elongated GSCs are enriched in the invasive and perivascular niches of primary tumour samples. (A-C) Representative microscopy images of peripheral GBM tissue expressing markers for the invasive niche (A and B): HuC/HuD (neurons) (A), IBA1 (microglia), MMP2 (invasive cancer cells); and markers for the perivascular niche (C): CD163 (perivascular macrophages) and PDGFRb (pericytes). Scale bars, 50 µm. (D) Quantitative distribution of GSC morphoclasses in invasive and perivascular niches vs tumor core (see Fig. 3E for the images of the core). Error bars, SEM. Two-way ANOVA with Tukey post-hoc test; Multipolar cells, ****, P<0,0001. Elongated cells, ***, P<0,001. N=3 patients (P12, P15, P17), 3 areas per patient section. **(E)** Four representative examples of assembloids composed of EdU+ core GBM organoid with a peripheral GBM organoid from 4 different patients (P07, P08, P11, P12). Note that the peripheral GBM organoids have been invaded by EdU+ cells from the core GBM organoids at different degrees depending on the patients. Scale bars, 500 µm.

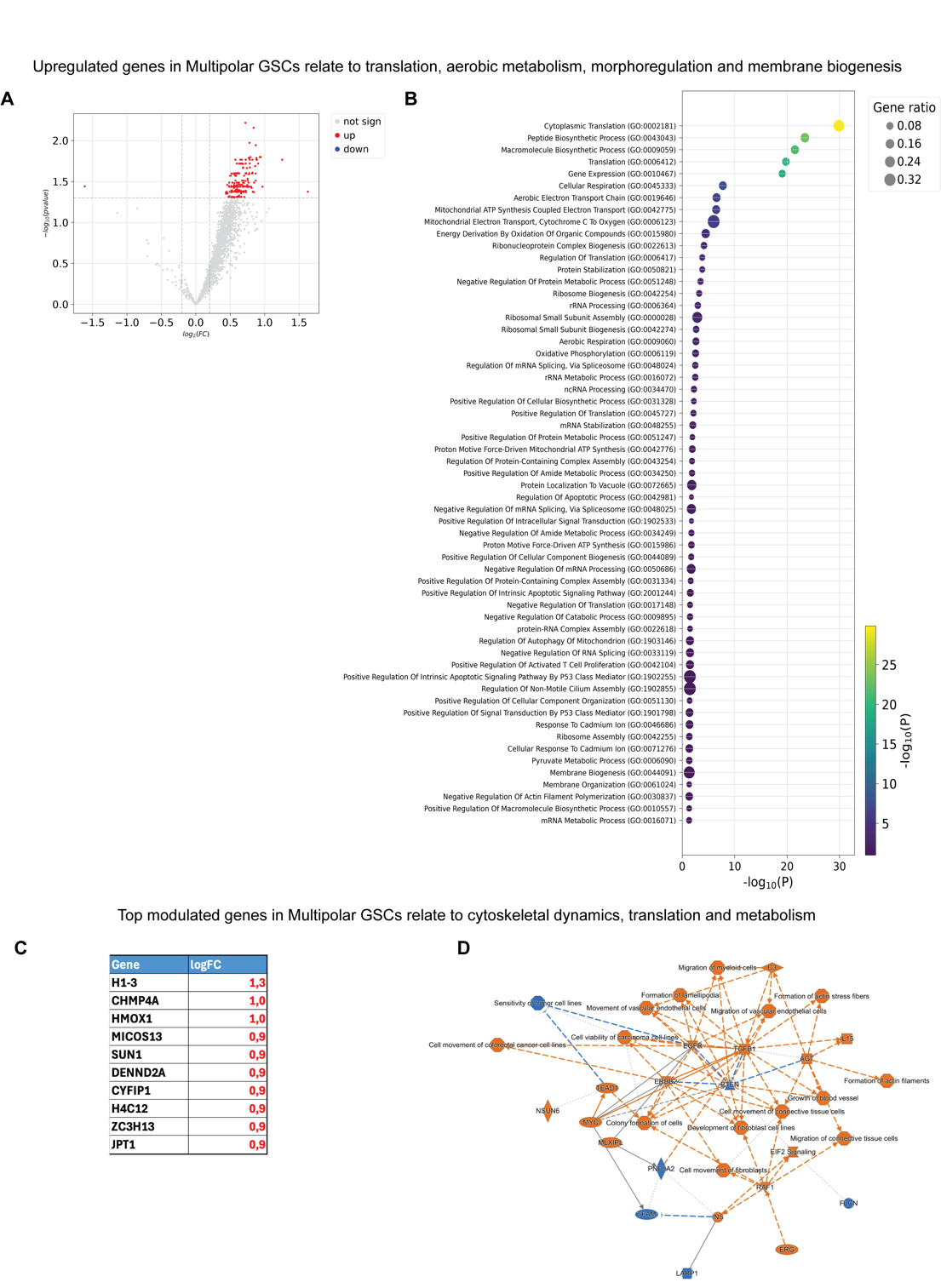

### **Figure S7. Multipolar GSCs are enriched in pathways related to protrusion growth, cytoskeletal dynamics and cell metabolism.**

**(A)** Differential expression analysis (DEA) of multipolar cells vs the rest. Volcano plot showing the results of a DEA, performed using PyDESeq2, comparing the expression profiles of AOIs with multipolar cells against the rest of the dataset. The x-axis represents Log-Fold Change (LFC), while the y-axis shows the negative log10 of the adjusted p-value. Dots highlighted in red are significantly up-regulated genes (LFC > 0 and adjusted p-value < 0.05), while dots highlighted in blue are significantly down-regulated genes (LFC < 0 and adjusted p-value < 0.05) **(B)** Upregulated pathways in multipolar GSCs. Significantly upregulated pathways identified through Over-Representation Analysis (ORA) using the *enrichr* function from *gseapy*. The analysis was performed on genes with significantly higher expression in ROIs containing elongated cells compared to the rest, selected based on a Log-Fold Change (LFC) > 0 and an adjusted p-value (corrected for multiple testing) < 0.05. In the plot, the x-axis position and dot color indicate the significance level of pathway enrichment, while dot size represents the proportion of differentially expressed genes associated with each GO term. **(C)** List of the top modulated genes with corresponding log fold change value observed in IPA Core Analysis for multipolar cells featuring genes involved in cytoskeleton dynamic (CYFP1, JPT1, SUN1) and cell metabolism (MICOS13, HMOX10) **(D)** Graphical summary providing an overview of the major biological themes observed in IPA Core Analysis for multipolar cells and their interrelations. It highlights the most significantly activated (orange) or inhibited (blue) canonical pathways (hourglass), diseases (cross), biological functions (octagon), and upstream regulators (rhombus for enzymes, square for cytokines, hexagon for translational regulators, oval for transcriptional regulators, inverted triangle for kinases, circle for others). The summary is constructed using machine learning techniques to prioritize and connect entities. Direct interactions are shown with solid lines, indirect interactions with dashed lines, and inferred edges with dotted lines.

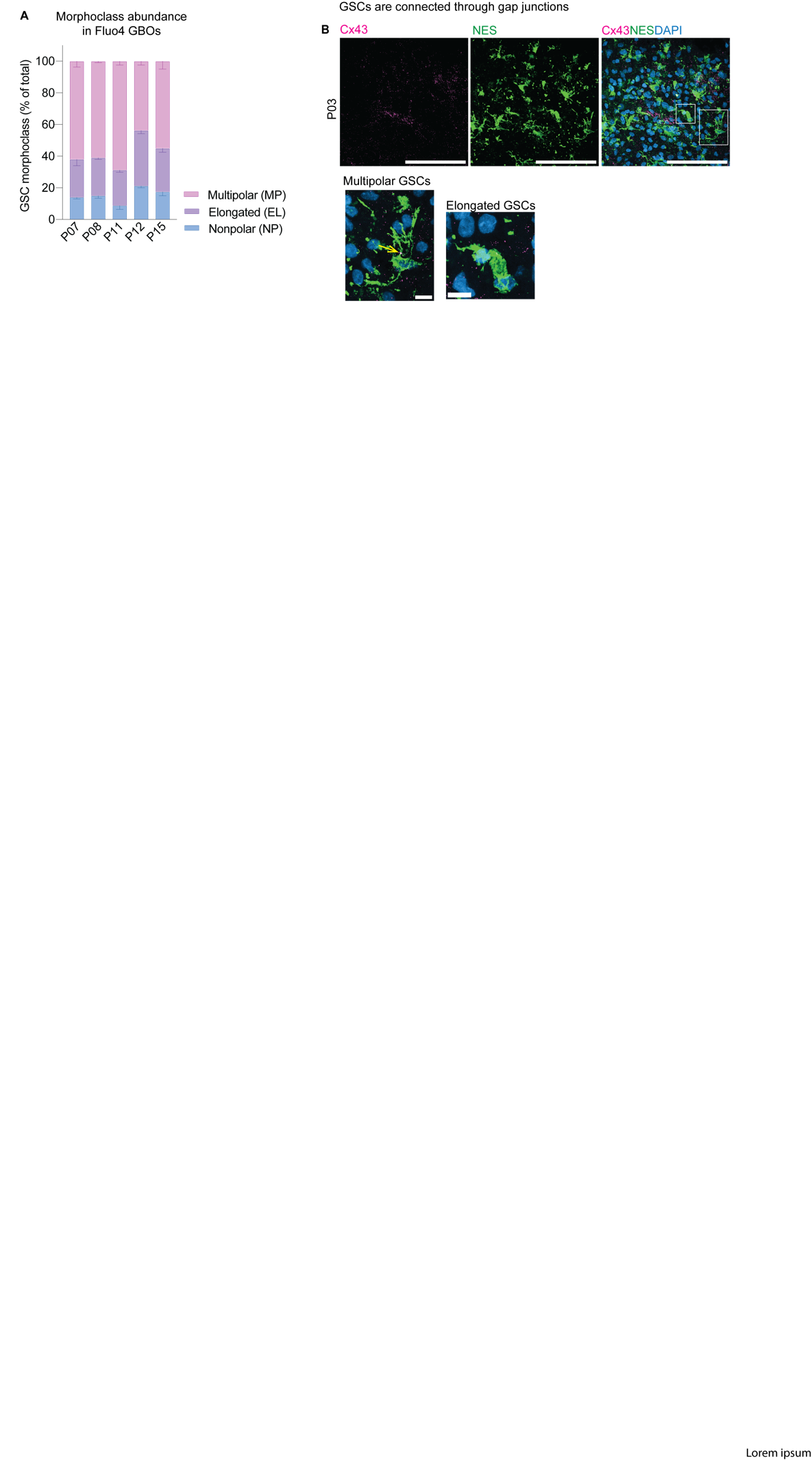

### **Figure S8. Multipolar cells are connected through gap junctions.**

**(A)** Quantitative distribution of the three morphoclasses in the patient GBM organoids used for live calcium imaging experiments. Error bars, SEM. N=5 patients (P07, P08, P11, P12, P15). **(B-F)** Representative microscopy images of patient-derived core GBM tissue stained with Connexin43 (magenta), Nestin (green) and DAPI (blue). (B) Note that multipolar cells are connected through Cx43+ gap junctions (yellow arrow). Scale bars, 100 µm (overview); 10 µm (close ups). N=4 patients (P03, P12, P15, P17).

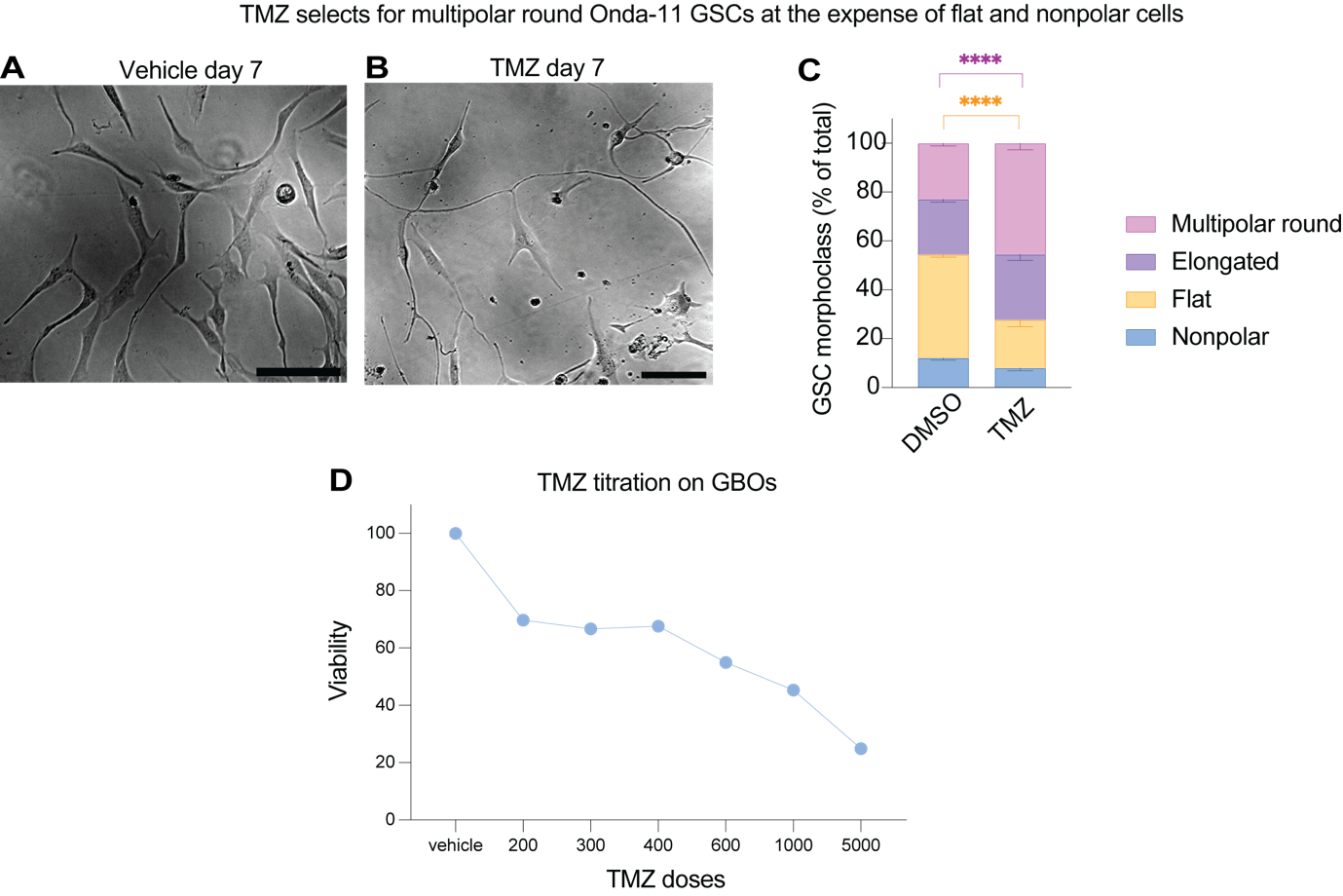

### **Figure S9. Multipolar GSCs are resistant to temozolomide.**

**(A-C)** Onda-11 GSCs are enriched in multipolar cells following chronic treatment with temozolomide (TMZ). (A-B) Representative bright field images of Onda-11 GSCs at day 7 following chronic administration of TMZ every 48h. Note that while in DMSO condition most cells are flat i.e. have a simple morphology (A), in the TMZ condition most cells are multipolar round (B). Scale bars, 30 µm. (C) Quantitative distribution of Onda-11 GSC morphoclasses at day 7 following metronomic 200 µM TMZ administration vs DMSO control from 2 independent experiments. ****, P<0,0001. **(B)** Titration curve of GBM organoids (GBO) treated with TMZ concentration ranging from 200 µM to 5000 µM. Y axes represent luminescence values normalized for the area of the organoids and expressed as normalization on the vehicle. N= 5 GBM organoids per condition, 1 patient (P09).

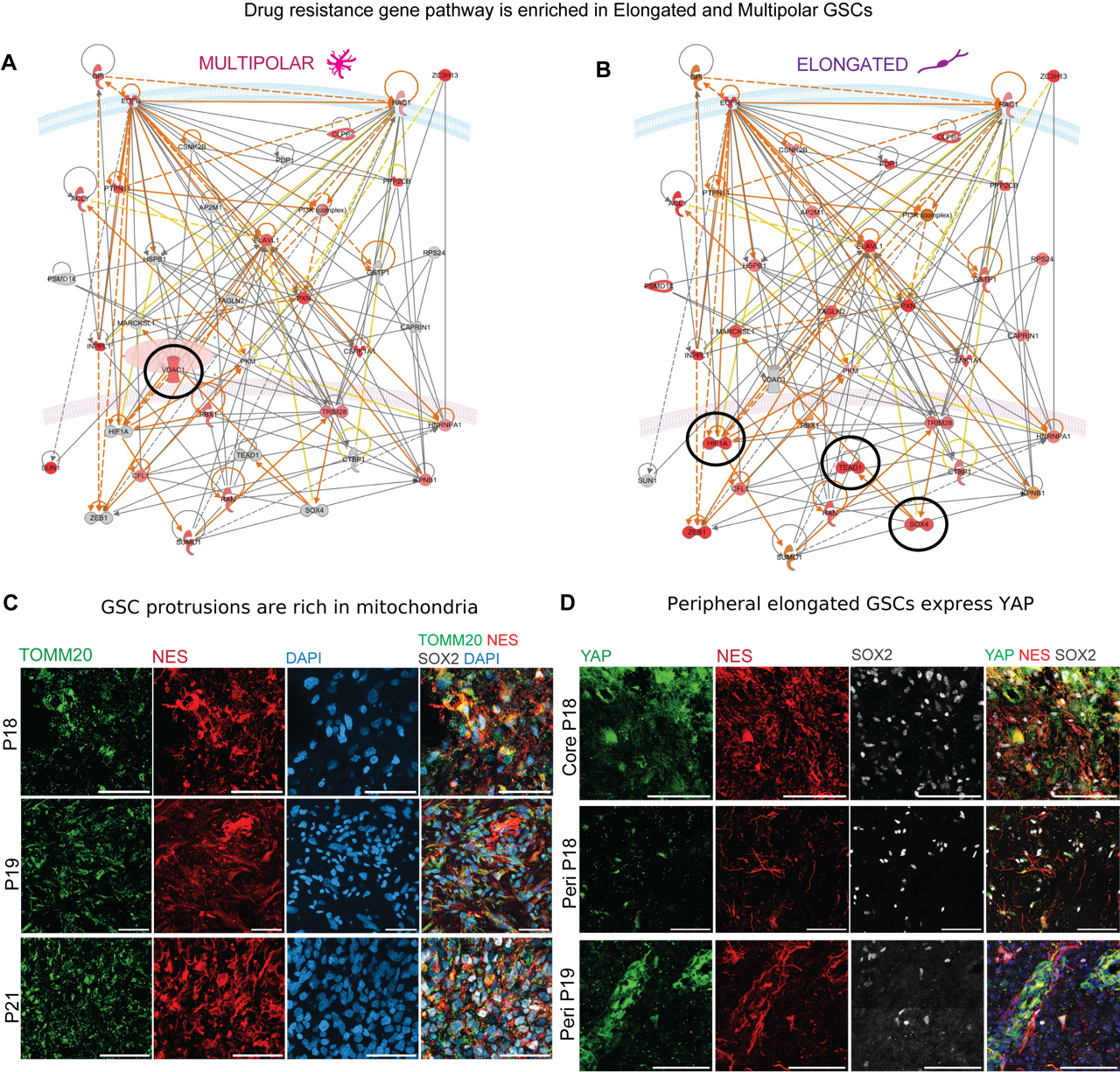

### **Figure S10. Multipolar and elongated GSCs have different signatures of therapy resistance.**

**(A, B)** Graphical representation of the molecular relationships between modulated genes related to tumor drug resistance function in multipolar cells (A) and elongated cells (B). Genes are depicted as nodes, and the biological relationship between two nodes is represented as an edge (line). All edges are supported by at least one reference from the literature, textbooks, or canonical information in the QIAGEN Knowledge Base. The intensity of the node color indicates the degree of up-regulation (red) or down-regulation (green). Nodes are displayed in various shapes representing the functional class of the gene product (diamond for enzymes, square for cytokines, hexagon for translational regulators, oval for transcriptional regulators, triangle for phosphatases, inverted triangle for kinases, rectangle for G-protein coupled receptors, circle for others, double circles for complexes). Highlighted in black are key genes specific to each of the two morphoclasses. **(C)** Representative microscopy images of GBM primary tissue (P18, P19, P21) containing Nes+ (red) SOX2+ (shown in merge as white) GSCs with mitochondria (TOMM20 immunofluorescence, green) in their protrusions. **(D)** GSCs express YAP. Note that in the tumor core (first row) YAP (green) is widely express while in periphery (lower two rows) YAP is present in the nucleus and cytoplasm of peripheral elongated Nes+ (red) Sox2+ (white) GSCs.

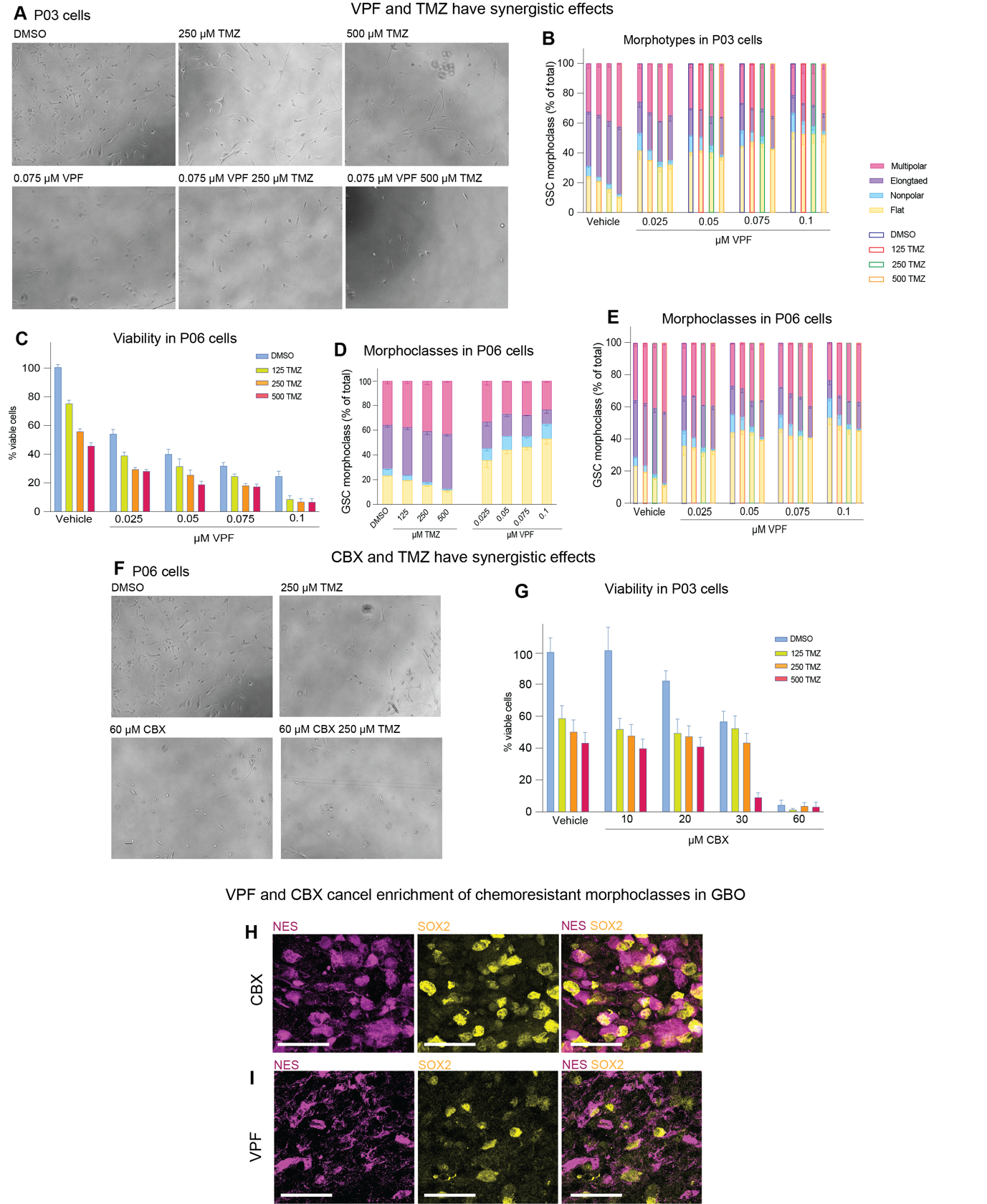

**Figure S11. GSCs have morphotype-specific vulnerabilities.**

**(A-E)** VPF has synergism with TMZ and depletes elongated GSCs. (A) Representative brightfield images of GSCs (P03, see Figure 5F and G for quantifications) treated with TMZ and VPF alone or in combination. (B) Morphotype distribution in GSCs treated with 0.025-0.1 µM VPF alone (blue stroke), or in combination with 125 µM TMZ (red stroke), 250 µM TMZ (green stroke), 500 µM TMZ (orange stroke). N=1 cell line (P03) in three biological replicates. Error bars, SEM. (C) The second GSC line (P06) corroborates the phenotypes observed in P03 GSC line. 0.025-0.1 µM VPF treatment alone (blue) or in combination with 125 µM TMZ (green), 250 µM TMZ (orange), 500 µM TMZ (red) for 96h. Measurements are normalized on DMSO control (100%). N=1 GSC line (P06) in three biological replicates. Error bars, SEM. (D) Morphotype distribution in GSCs (P06) treated with 0.025-0.1 µM VPF and 125-500 µM TMZ. (E) Morphotype distribution in GSCs treated with 0.025-0.1 µM VPF alone (blue stroke), or in combination with 125 µM TMZ (red stroke), 250 µM TMZ (green stroke), 500 µM TMZ (orange stroke). N=1 cell line (P06) in three biological replicates. Error bars, SEM. **(F, G)** CBX has synergism with TMZ. (F) Representative brightfield microscopy images of GSCs (P06, see Figure 5H and I for quantifications) treated with TMZ and CBX. Note that CBX depletes multipolar GSCs and enriches for elongated GSCs. (G) The second GSC line (P03) corroborates the phenotypes observed in the P06 GSC line. GSC viability after 96h of 10-60 µM CBX treatment alone (blue) or in combination with 125 µM TMZ (green), 250 µM TMZ (orange), 500 µM TMZ (red). Measurements are normalized on DMSO control (100%). N=1 GSC line (P03) in three biological replicates. Error bars, SEM. **(H, I)** VPF and CBX cancel the enrichement of chemoresistant morphotypes in GBOs. Representative microscopy images of GBOs (P24) treated for 10 days with 60 µM CBX (H) and 0.075 µM VPF (I) and stained for nestin (magenta) and SOX2 (yellow) to identify GSCs. Note the enrichement of nonpolar cells in both conditions. See also Figure 5K-N.

**Tables**

**Table S1. Reagents table.** For each reagent including cell lines, antibodies, chemicals and softwares, the source and identifier are noted.

**Table S2. Patients table.** For each patient sample, its code, age, sex, histological diagnosis, collected tumour areas and experiments performed on it are shown.

**Movies**

**Movie S1.** Representative example of Fluo4 calcium imaging movie on patient P11. 100 frames per second (fpm). Frame width, 389 µm.

**Movie S2.** Representative example of Fluo4 calcium imaging movie on patient P13. 100 frames per second (fpm). Frame width, 389 µm.

**Movie S3.** Representative example of Fluo4 calcium imaging movie on patient P14. 100 frames per second (fpm). Frame width, 389 µm.

**Movie S4.** Representative example of Fluo4 calcium imaging movie on patient P14 following 15 minutes treatment with Carbenoxolone (CBX). 100 frames per second (fpm). Frame width, 389 µm.

**References present in the Supplemental figure legends**

1	Garofano, L. *et al.* Pathway-based classification of glioblastoma uncovers a mitochondrial subtype with therapeutic vulnerabilities. *Nat Cancer* **2**, 141-156, doi:10.1038/s43018-020-00159-4 (2021).
